# Progressive neuronal network reorganisation in glioblastoma drives pathological activity *in vitro*

**DOI:** 10.64898/2026.07.16.738125

**Authors:** Giulia Amos, Luc Jordi, Karan Ahuja, Alexandra Gerber, Gregor Hutter, János Vörös, Christina M. Tringides, Vaiva Vasiliauskaitė

## Abstract

Glioblastoma (GBM) is the most aggressive primary brain tumour and is frequently accompanied by severe neurological symptoms, including epilepsy and cognitive impairment. Neurological symptoms often persist after surgical resection, indicating that GBM induces durable and self-sustaining changes in the surrounding neuronal networks. However, the mechanisms by which GBM reshapes network structure and function in the tumour periphery remain poorly understood. We present a compartmentalised *in vitro* platform enabling long-term coculture of iPSC-derived neurons and primary GBM cells to investigate these changes. Placed on high-density microelectrode arrays, the platform permits longitudinal electrophysiological recordings at single-neuron resolution. Using effective network inference, we find that GBM drives a reproducible structural progression: first toward a hyperconnected, hub-dominated architecture, then a collapse of community structure accompanied by a widespread neuron loss. This evolving structure shapes population dynamics, constraining features such as network burst rate and instantaneous synchrony. The reorganisation also carries computational consequences: signal propagation becomes progressively redundant and synergistic rather than unique. As a result, neurons lose the capacity to encode distinct input combinations independently, and the repertoire of accessible network states contracts. Together, these findings reframe GBM as a driver of neuronal network reorganisation rather than uniform hyperexcitability, and establish a compartmentalised, single-neuron-resolution platform for the longitudinal observation, dissection, and ultimately targeting of the network processes that underlie disease progression.

## Introduction

Glioblastoma (GBM) is the most aggressive primary brain tumour, with a five-year survival rate of 5% (1, 2). Despite extensive research efforts, therapeutic advances have been limited and patient prognosis has remained largely unchanged over the past two decades (3). Historically, GBM research has focused primarily on tumour-intrinsic properties such as molecular characteristics, genetic alterations, and cellular composition (4–6). However, increasing evidence suggests that understanding GBM progression requires moving beyond a tumour-centric perspective towards studying GBM as a disease of the full central nervous system (7, 8). This shift recognises that tumour progression is strongly shaped by interactions between GBM cells and the surrounding neuronal environment.

Rather than existing in isolation, GBM actively integrates into the brain through bidirectional interactions. GBM cells have been shown to form functional synapses with neurons, exploit glutamatergic signalling pathways that promote proliferation, and establish extensive cellular networks through tumour microtubes (9–13). These studies have provided important mechanistic insights into neuron–GBM interactions at the cellular and microscale level. Emerging evidence suggests that GBM-induced cellular interactions may extend beyond local circuit alterations and contribute to widespread changes in network organisation, with recent neuroimaging studies reporting altered structural and functional connectivity at the whole-brain level in patients with GBM (14– 17). However, these observations provide only a coarsegrained view of network disruption and limited insight into how cellular-scale neuron–GBM interactions progressively remodel mesoscale neuronal networks at and beyond the tumour periphery.

Network neuroscience provides a framework for addressing these questions by representing neuronal elements — whether individual neurons or whole brain regions — as nodes and their interactions as edges within a graph (18, 19). Applied across a range of neurological disorders, this approach has revealed characteristic patterns of network remodelling, including hub disruption (20), altered modular structure (21), and changes in information flow (22). Importantly, network structure is not merely descriptive: connectivity constrains how activity propagates through a network and therefore shapes the dynamical and computational states that the network can support (23). Changes in connectivity can consequently alter not only patterns of neuronal activity, but also how information is transmitted and integrated across neuronal populations to support cognition (24).

This structure–dynamics perspective is particularly relevant for GBM. A substantial proportion of GBM patients develop epilepsy (25), and many experience cognitive impairment over the course of the disease (26, 27). Notably, these symptoms often persist after surgical resection (28, 29), suggesting that tumour-associated network alterations are self-sustained and not reversed by the removal of the bulk tumour alone. At the whole-brain scale, gliomas preferentially arise in highly connected hub regions of the human connectome and are associated with large-scale functional changes (30, 31). Whether GBM’s progressive remodelling of neuronal connectivity systematically reshapes mesoscale network architecture, and whether this reorganisation can explain the onset and persistence of clinical symptoms, remains unknown.

One reason for this gap is the difficulty of tracking mesoscale neuronal networks during GBM progression *in vivo*. Experimental access is limited, measurements are subject to multiple confounding factors, and current approaches generally lack the resolution required to follow network reorganisation at single-neuron scale over extended periods. *In vitro* systems provide a complementary approach. In particular, compartmentalised platforms incorporating human induced pluripotent stem cell (iPSC)-derived neuronal networks enable controlled investigation of disease processes while maintaining direct human relevance (32, 33). Combined with high-density microelectrode arrays (HD-MEA), these systems permit stable long-term recordings at high spatiotemporal resolution, enabling longitudinal analysis of network remodelling (34, 35).

Here, we present a compartmentalised *in vitro* platform designed to model the tumour periphery and enable longitudinal investigation of neuron–GBM interactions over multiple months. Because GBM comprises transcriptional subtypes with distinct genetic drivers and phenotypes (36), we consider both the classical and mesenchymal molecular classes (37–39) to test whether they converge on a distinct or shared pattern of network reorganisation. Generally, the classical subtype is characterised by *EGFR* amplification and high proliferative activity, whereas the mesenchymal subtype adopts a more invasive, inflammation-associated phenotype (40). Using multivariate transfer entropy (mTE), we infer directed effective connectivity networks and track their structural properties from early infiltration through late-stage neuronal death. We show that GBM drives a progressive and reproducible reorganisation of network topology characterised by the emergence of broadcasting hubs and collapse of community structure. This structural reorganisation constrains the dynamical regimes accessible to the network and reshapes how information is transmitted and integrated across neuronal populations. Specifically, GBM shifts information processing from unique transmission towards redundant and synergistic integrations. Together, these findings establish a testable mechanistic framework connecting mesoscale network reorganisation to functional and computational changes.

## Results

### Compartmentalised platform enables longitudinal monitoring of neuron–GBM interactions

We developed a three-compartment polydimethylsiloxane (PDMS) microstructure platform that spatially segregates iPSC-derived neurons from patient-derived primary GBM cells to model the tumour periphery *in vitro* (Fig. 1 a,b). Each structure consists of two neuronal seeding wells connected to a lateral GBM well via ten microchannels. The microchannels incorporate axon-guidance features and somatic constrictions (1.5 *µ*m) that direct neurite outgrowth while restricting neuronal cell bodies to the wells, allowing axons to contact GBM cells at a defined distance from the soma. The compartmentalisation prevented the GBM cells from overgrowing the neuronal population, and preserved neuronal viability over the co-culture timescales required to study network-level reorganisation.

**Fig. 1.**
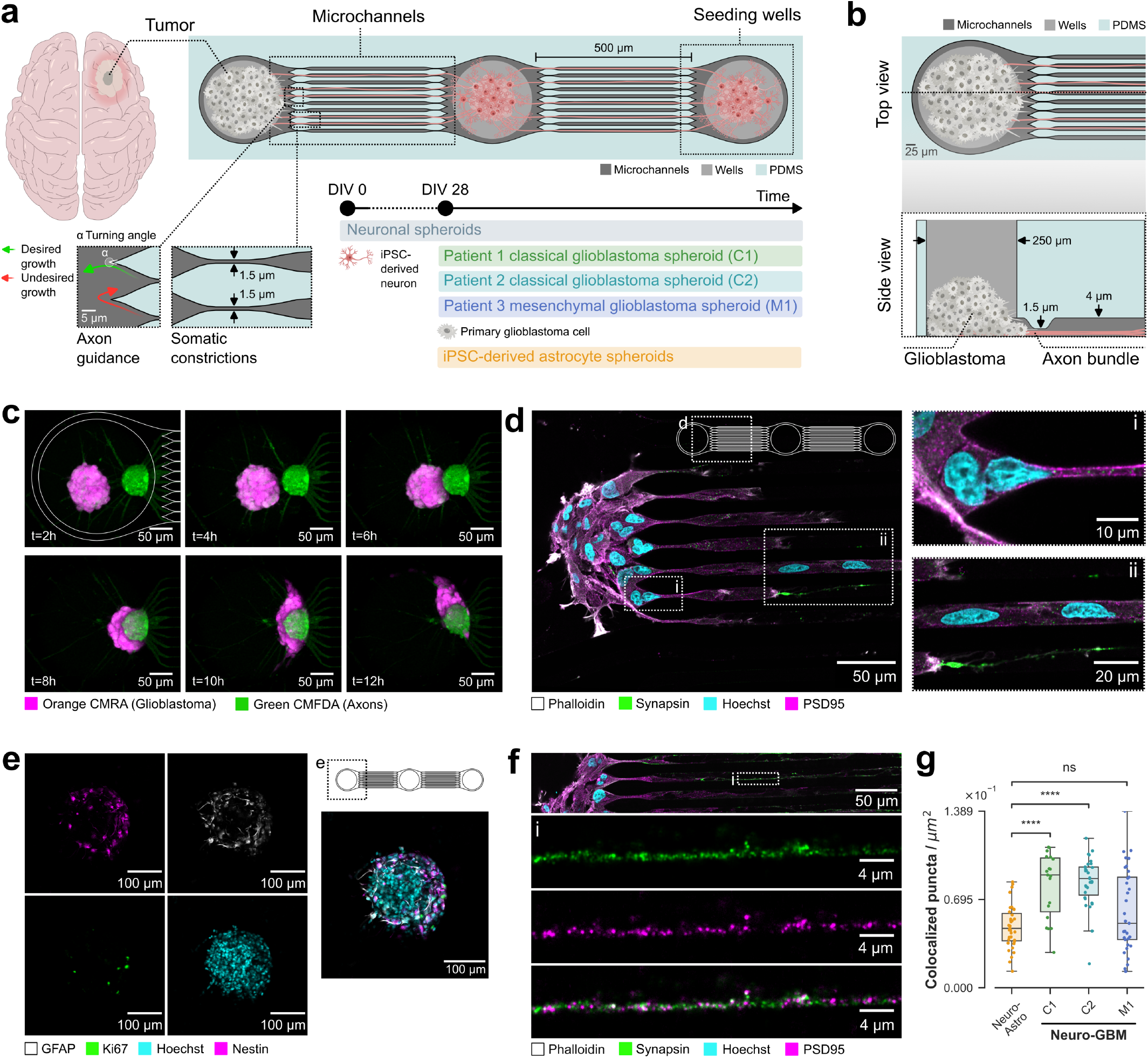
Compartmentalised microfluidic platform for long-term monitoring of glioblastoma (GBM) invasion and tumour–neuron interactions *in vitro*. **a** Schematic of the microfluidic device comprising three seeding wells interconnected by microchannels. 200 iPSC-derived neurons were seeded as spheroids into two of the three wells and cultured for 28 days prior to the addition of 100-cell patient-derived primary GBM spheroids or control astrocyte spheroids into the third compartment. Insets show axon-guiding structures and somatic constrictions that restrict neuronal cell soma migration while permitting axonal growth. **b** Cross-section schematic of the same device, illustrating the spatial relationship between axons and the GBM spheroid. **c** Representative timelapse images showing a GBM spheroid (magenta) approaching an axonal bundle (green) at the microchannel entry within the first hours following co-culture initiation. **d** GBM cells show signs of nuclear softening, enabling them to deform and migrate through the narrow microchannels (DIV 35). **e** Immunocytochemistry confirms retention of a cancerous phenotype in co-culture. **f–g** Quantification of co-localised synaptic puncta (Synapsin I /PSD-95) within microchannels at DIV 35. Both classical Neuro–GBM conditions (C1 and C2) exhibited a significant increase in puncta density relative to the Neuro–Astrocyte control (Kruskal–Wallis across conditions followed by Dunn’s post-hoc, ns, not significant, p > 0.05; *p < 0.05; **p < 0.01; ***p < 0.001; ****p < 0.0001). Boxplots show median and inter-quartile range; points show individual 10-channel areas.

iPSC-derived excitatory neurons were seeded as 200-neuron spheroids into two compartments and cultured for 28 days *in vitro* (DIV) to allow the network to mature. At DIV 28, 100-cell spheroids of primary GBM cells were introduced into the lateral compartment using three patient-derived lines: two classical subtypes (C1, “Neuro-GBM C1” and C2, “Neuro-GBM C2”) and one mesenchymal subtype (M1, “Neuro-GBM M1”). iPSC-derived astrocytes (“Neuro-Astro”) and neuron-only monocultures (“Neuro”) served as non-tumour and negative controls, respectively. Characterisation of axonal growth, tumour proliferation and astrocyte markers in the microstructures can be found in Supplementary Figure S1.

Before using the platform to study network-level reorganisation, we confirmed that GBM cells retained their characteristic behaviour in the microstructure. Within hours of seeding, GBM cells migrated toward the axon bundle, established direct contact, and wrapped around it (Fig. 1 c; Supplementary videos show interaction over first 12 h). Over time, cells extended processes into the microchannels and displayed nuclear morphology consistent with nuclear deformation (Fig. 1 d) (41). Ki67, Nestin, and GFAP immunostaining confirmed that GBM cells maintained a proliferative, cancerous phenotype in co-culture (Fig. 1 e). Synaptic puncta density, quantified as colocalised Synapsin/PSD95 puncta along axon bundles, was significantly higher in neurons co-cultured with classical GBM lines C1 and C2 than in the astrocyte control at DIV 35 (Kruskal-Wallis with posthoc Dunn test, *p* < 0.0001; Fig. 1 f,g), consistent with previously reported GBM-driven synaptogenesis (9). No significant difference was observed for the mesenchymal line M1. Collectively, these observations confirm that GBM cells actively interact with neurons in the microstructure. Because the microstructure physically separates the tumour from the neuronal network at seeding, tumour infiltration occurs gradually, enabling non-invasive recordings of the same cultures over weeks. The compartmentalised platform therefore allows longitudinal observation of neuronal network responses to spatially confined neuron–GBM interactions.

### HD-MEA recordings combined with multivariate transfer entropy resolve directed effective connectivity in co-cultures

To extract network-level information from the co-cultures, we recorded electrophysiological activity using a HD-MEA onto which the PDMS microstructure was aligned. The setup fits nine identical networks onto a single chip, enabling parallel recordings across networks (Fig. 2 a,b). The microchannels measure 10 *µ*m in width with an inter-channel spacing of 12 *µ*m, ensuring that each electrode records from a single channel (electrode pitch: 17.5 *µ*m). The channel height of 4 *µ*m confines the axon bundle in close proximity to the electrodes, yielding high signal-to-noise ratio recordings with spike amplitudes reaching above 400 *µ*V (Fig. 2 c,d).

**Fig. 2.**
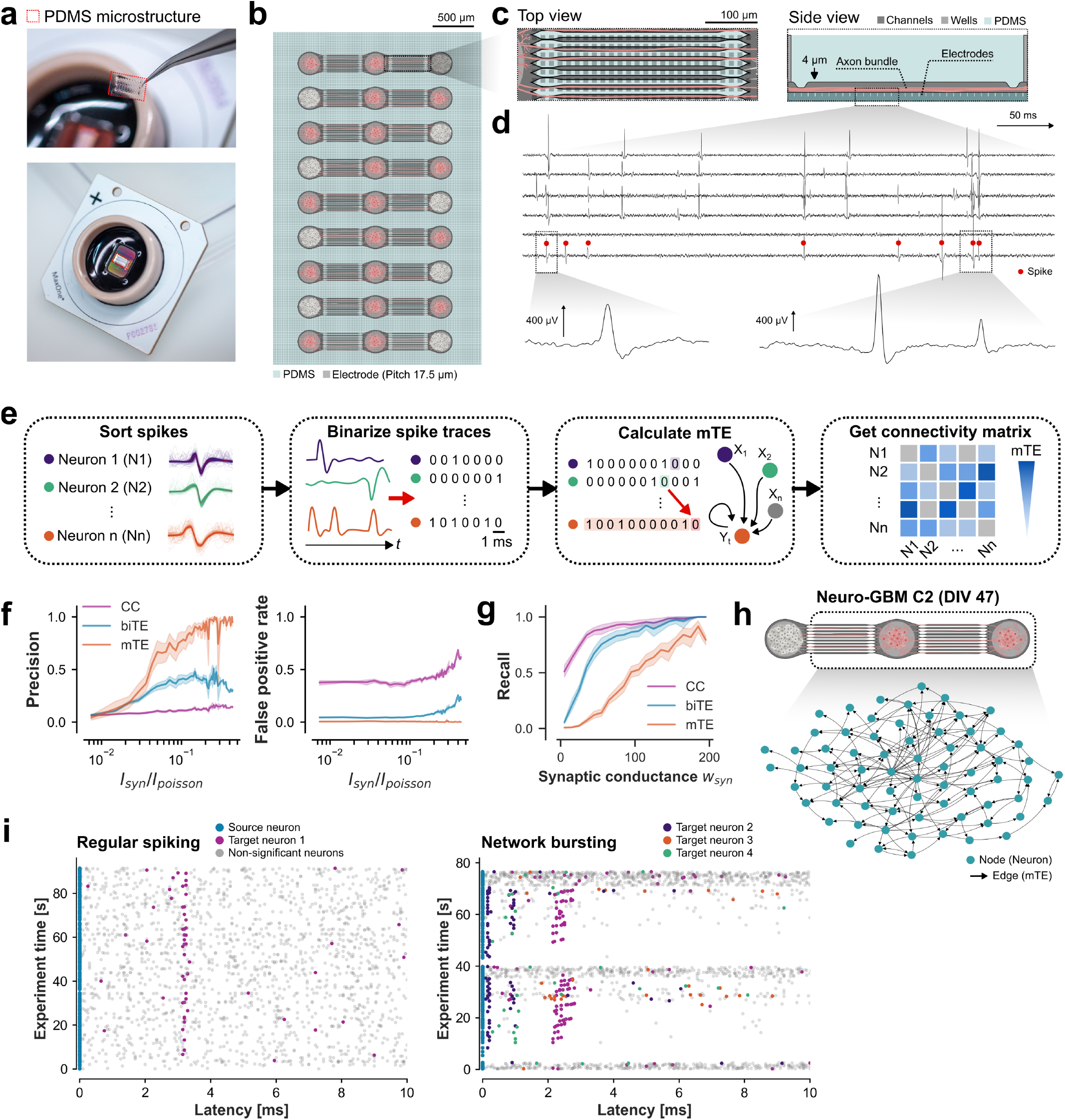
Multivariate transfer entropy enables inference of effective connectivity from extracellular spike recordings. **a** Photograph showing alignment of the microfluidic platform with the sensing area of a CMOS microelectrode array (MEA). **b** Schematic of parallel recording from up to nine spatially separated networks on a single MEA. **c** Functional signals are acquired from electrodes positioned beneath the microchannels of each network. Cross-section view illustrates the close contact between axon bundles and electrodes within the channels. **d** Confinement within the narrow microchannels amplifies extracellular signals, yielding spike traces with high signal-to-noise ratios. Individual spikes are indicated with red dots for the last voltage trace. **e** Analysis pipeline for inference of effective connectivity: spike trains are first binarised, then effective connectivity is inferred using multivariate transfer entropy (mTE). **f** Precision and false-positive rate of three inference methods (cross-correlation, CC; bivariate transfer entropy, biTE; multivariate transfer entropy, mTE) as a function of synaptic to Poisson current ratio *I*_*syn*_*/I*_*poisson*_. mTE achieves the highest precision and a nearzero false-positive rate across the full range. **g** Recall as a function of synaptic conductance *w*_*syn*_. All methods recover strong connections reliably; mTE’s recall is lowest for weak connections but rises steadily with *w*_*syn*_, indicating that the edges it misses are predominantly the weakest ones. **h** Representative effective connectivity network reconstructed from functional recordings of a Neuro–GBM C2 co-culture at DIV 47. **i** Spike-time-triggered raster plots demonstrating accurate identification of synaptic pairs under both tonic firing and network bursting regimes.

We implemented a multivariate transfer entropy (mTE) (42) pipeline (Fig. 2 e) to infer directed effective network connectivity from the electrophysiological recordings. Spikes were first sorted from raw traces, binarised and binned at 1 ms resolution. mTE was then computed between all neuron pairs to yield a weighted, directed connectivity matrix whose entries (edges) quantify the unique directed influence of a source neuron on a target neuron (see “Inference of effective connectivity” in Methods). Unlike pairwise methods such as cross-correlation (CC) or bivariate TE (biTE), which report all statistical dependencies between pairs regardless of whether they are direct or mediated by other neurons, mTE identifies a minimal directed model of the network by requiring each inferred connection to provide statistically significant information about the target that is not already explained by other sources (18). This iterative conditioning eliminates redundant connections and captures synergistic interactions (42, 43), producing sparser and more accurate network representations. As a result, effective networks derived from mTE are better suited to the computation of graph-theoretic measures than functional networks.

To benchmark mTE against standard pairwise methods, CC and biTE, we used Adaptive Exponential Leaky Integrateand-Fire (AdExLIF) network simulations with a known ground-truth graph (see “Validation of effective connectivity inference” in Methods and Supplementary Figure S2) (44). Across a range of synaptic currents, mTE achieved substantially higher precision and a much lower false positive rate than both CC and biTE, at the cost of reduced recall (Fig. 2 f,g). The lower recall reflects a known limitation of mTE: weak connections convey insufficient unique information to pass the conditional significance threshold, and are therefore more likely to go undetected (42). In a neuroscientific context, this trade-off is favourable: false positives are considerably more detrimental to network-level interpretations than missed weak connections (45). Importantly, mTE maintains high precision even when network activity is highly correlated in contrast to the pairwise methods which severely overestimate connectivity due to redundant shared activity (Fig. 2 f and S2). Since GBM-driven hyperexcitability is expected to increase network-wide correlations in the co-culture data, this is an important consideration.

Applied to empirical data, the pipeline produced welldefined directed network graphs from co-culture recordings (Fig. 2 h), and reliably identified directed neuronal pairs in both regular spiking and network bursting regimes (Fig. 2 i).

### Glioblastoma drives network reorganisation toward broadcasting hub dominance and community collapse

We next asked whether neuron–GBM interactions *in vitro* produce measurable reorganisation of effective network structure, from early infiltration to the late-stage loss of neurons. To address this, we quantified both local and global properties of effective networks recorded longitudinally from DIV 26 to DIV 60. (Fig. 3 a,b), with DIV 26–28 serving as baseline before GBM or astrocyte addition. All extracted features are described in Supplementary Table 4.

**Fig. 3.**
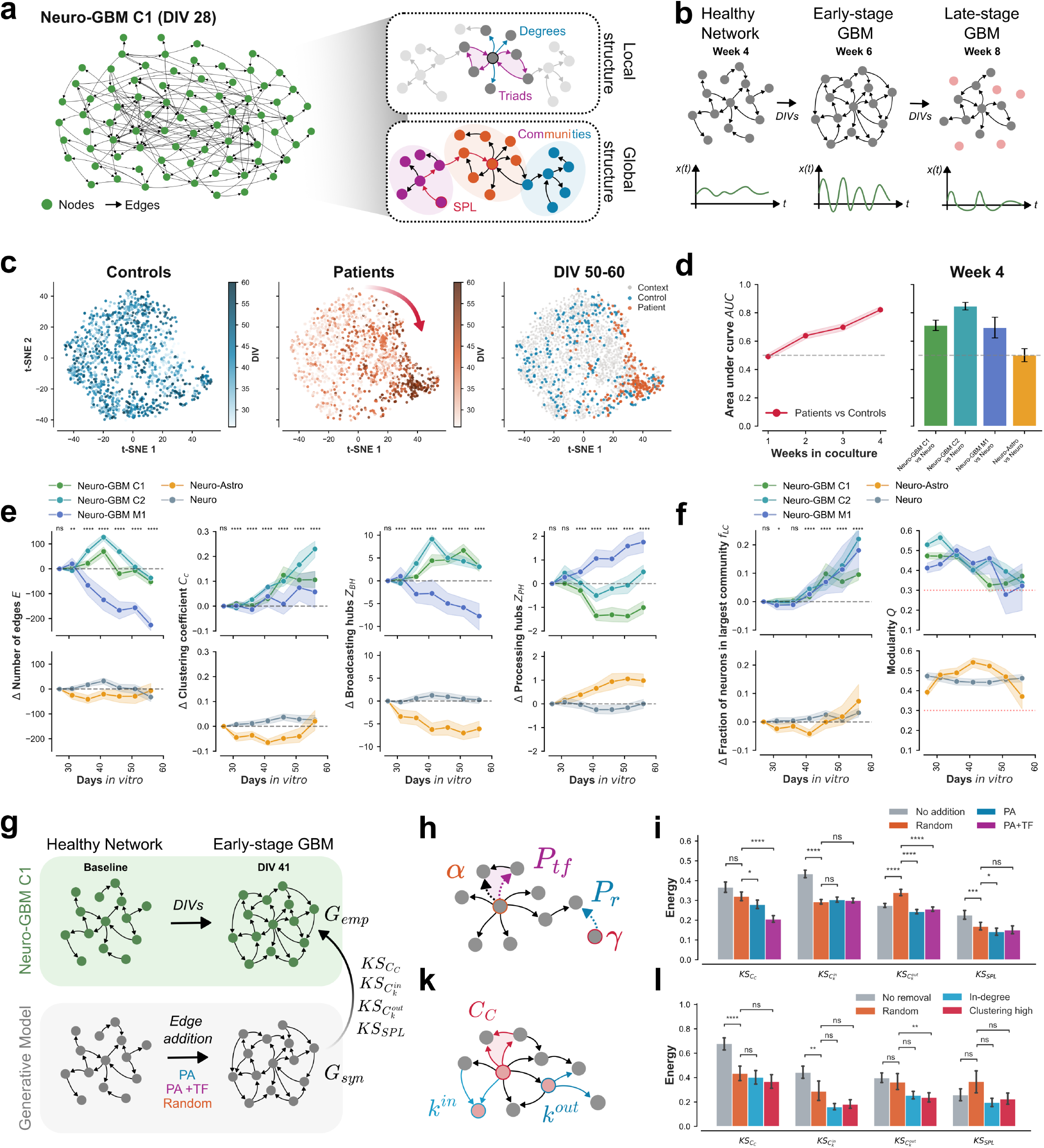
Glioblastoma (GBM) infiltration leads to emergence of strong broadcasting hubs and community collapse. **a** Local (degree distributions, triad formation) and global (communities, shortest path length) network features are calculated from the effective connectivity graphs. **b** Using these metrics, we quantify the continuous reorganisation in network topology during GBM progression, divisible into an early growth phase and a late node-loss phase. **c** t-SNE embeddings of topological features reveal no pattern in control conditions (Neuro-Astro and Neuro, left), but a clear trajectory in patient conditions (Neuro-GBM C1, C2 and M1, middle). At DIV 50-60, the patient conditions show a clear cluster. Each point represents one network. **d** Leave-one-chip-out area under curve (AUC) of trained regularised logistic regression per bin shows topological divergence across conditions and coculture weeks. Lineplots and Barplots represent mean ± standard deviation. **e** Longitudinal change relative to baseline in the number of edges, clustering coefficient, broadcasting and processing hub z-score over days *in vitro*. Classical GBM promotes edge formation and broadcasting hub emergence, while mesenchymal GBM triggers rapid connectivity loss; clustering coefficients increase in both cases. Lineplots represent mean ± 95% CI; Data binned in 5 DIVs. **f** Community structure initially expands before collapsing as modularity falls below 0.3 (red line), signalling loss of network segregation. **g–i** Generative models were fitted to empirical networks using an energy metric *E*. Parameters (*α, γ, P*_*tf*_ , *P*_*r*_) were optimised via Bayesian search with stratified cross-validation. Preferential attachment with triad formation (PA+TF) best captured early-stage connectivity changes. Barplots represent mean; error bars 95 % CI. **k–l** For the late-stage decline, node removal mechanisms (random, degree-based, clustering-based) were evaluated on top of fixed PA+TF parameters, with selection strength (*δ*) and removal probability (*P*_*d*_) optimised per mechanism. Stars indicate, within each panel, a significant difference between conditions (Kruskal–Wallis across conditions followed by Dunn’s post-hoc with Benjamini-Hochberg correction, ns, not significant, *p >* 0.05; \**p <* 0.05; \*\**p <* 0.01; \*\*\**p <* 0.001; \*\*\*\**p <* 0.0001).

Projecting all network-level structural features into a lowdimensional space using t-SNE (46) revealed that network structure of GBM co-cultures follow a characteristic trajectory over time (Fig. 3 c). By DIV 50–60, networks from GBM co-cultures clustered in a distinct region of feature space, whereas neuron-only and neuron-astrocyte controls remained broadly distributed, indicating that GBM induces a convergent structural phenotype not shared by controls. A logistic regression classifier distinguished GBM co-cultures from controls with progressively higher accuracy, reaching an area under the curve (AUC, where 0.5 = chance and 1.0 = perfect discrimination) of 0.82 by week 4 in co-culture, with GBM networks diverging more strongly from neuron-only controls than neuron-astrocyte co-cultures. (Fig. 3 d).

At the local level, classical GBM co-cultures showed an initial increase in the average number of pairwise connections, followed by a progressive decline (Fig. 3 e). Clustering coefficient increased over the early co-culture period, accompanied by a rise in broadcasting hubs and a reduction in processing hubs (Fig. 3 e, as reflected in shifts of the out- and in-degree centrality distributions; degree and shortest path length distributions shown in Fig. S3 a–f). At the global level, the fraction of neurons belonging to the largest community initially increased before community structure collapsed at late time points, with modularity falling below 0.3, which is the threshold below which modular structure is considered absent (47) (Fig. 3 f). This collapse occurred in 56 % of networks for Neuro-GBM M1, 34 % of networks for Neuro-GBM C2 and 25 % of networks for Neuro-GBM C1 by DIV 56. The mesenchymal line Neuro-GBM M1 showed the opposite trend for several metrics: the combination of a more aggressive phenotype (40, 48) and an early loss of pairwise connections suggests that neuronal death proceeds more rapidly in this subtype. Neuron-only and neuron-astrocyte controls remained comparatively stable across all metrics.

We used generative modelling framework (49) (see “Generative Models” in Methods) to mechanistically account for the observed reorganisation in classical GBM co-cultures. Generative models grow networks by iteratively adding edges according to a specified wiring rule, allowing us to test whether simple, biologically interpretable mechanisms are sufficient to reproduce the empirical network topology. Models were compared using an energy metric 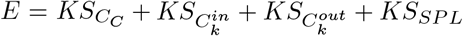 (adapted from (50)), where each term is a Kolmogorov–Smirnov statistic quantifying the discrepancy between simulated and empirical distributions, and lower energy indicates greater overall similarity to the observed network topology (Fig. 3 g). During the early stage (DIV 28–41), when edges are predominantly added, both preferential attachment (PA) and preferential attachment with triad formation (PA+TF) outperformed random attachment (Fig. 3 h,i), indicating that new connections preferentially target already well-connected neurons. PA+TF fit best by a margin of 0.21 (18.8% reduction in energy) compared to random, additionally reproducing the rise in clustering coefficient through its triad-closing mechanism.

The late stage (DIV 41–58) is instead dominated by neuron loss. Starting from the best-fitting PA+TF parameters, we compared node removal targeted by in-degree, out-degree, or clustering coefficient against non-selective random removal (Fig. 3 k). Removal of high-clustering nodes performed only marginally better than the alternatives (only significant comparison: random vs. high-clustering removal, p = 0.02, Dunn post-hoc test), suggesting that late-stage neuron loss is largely non-selective (Fig. 3 l). Example empirical network trajectories and best-fitting parameter values are provided in the Supplementary Material (Fig. S3 g–j).

These results suggest that GBM drives a reproducible and progressive restructuring of neuronal network topology, with an initial transition toward a hyperconnected, broadcasting hub-dominated architecture, followed by a collapse of community structure and widespread neuron loss. That this trajectory can be partially recapitulated by preferential attachment followed by node loss suggests the reorganisation is not random, but governed by activity-dependent wiring rules.

### Global effective network structure predicts population dynamics

Over disease development, GBM co-cultures diverged from controls in their population dynamics as well as amongst each other (Fig. 4 a,b). In classical GBM, activity became progressively more synchronous and network burstdominated, with pairwise synchrony (rCC), network burst rate (NBR), and network burst duration (NBD) all rising relative to baseline. The mesenchymal GBM followed the opposite trajectory: network burst rate and duration fell sharply, and population activity had largely collapsed by DIV 47 (Fig. 4 a). Neuron-only controls remained comparatively stable across all three measures, whereas neuron-astrocyte cocultures showed a modest decline (Fig. 4 b). These dynamical changes parallel the structural reorganisation described above. Effective connectivity and population dynamics are derived from the same spike trains but capture the data at different timescales: mTE identifies the minimal directed network explaining statistical dependencies between individual neurons at the millisecond scale of synaptic transmission, whereas network bursts are a collective phenomenon unfolding over seconds. The two are therefore not redundant by construction, which raises a natural question: does the inferred network architecture carry predictive information about population dynamics beyond what is already implicit in the spike trains?

**Fig. 4.**
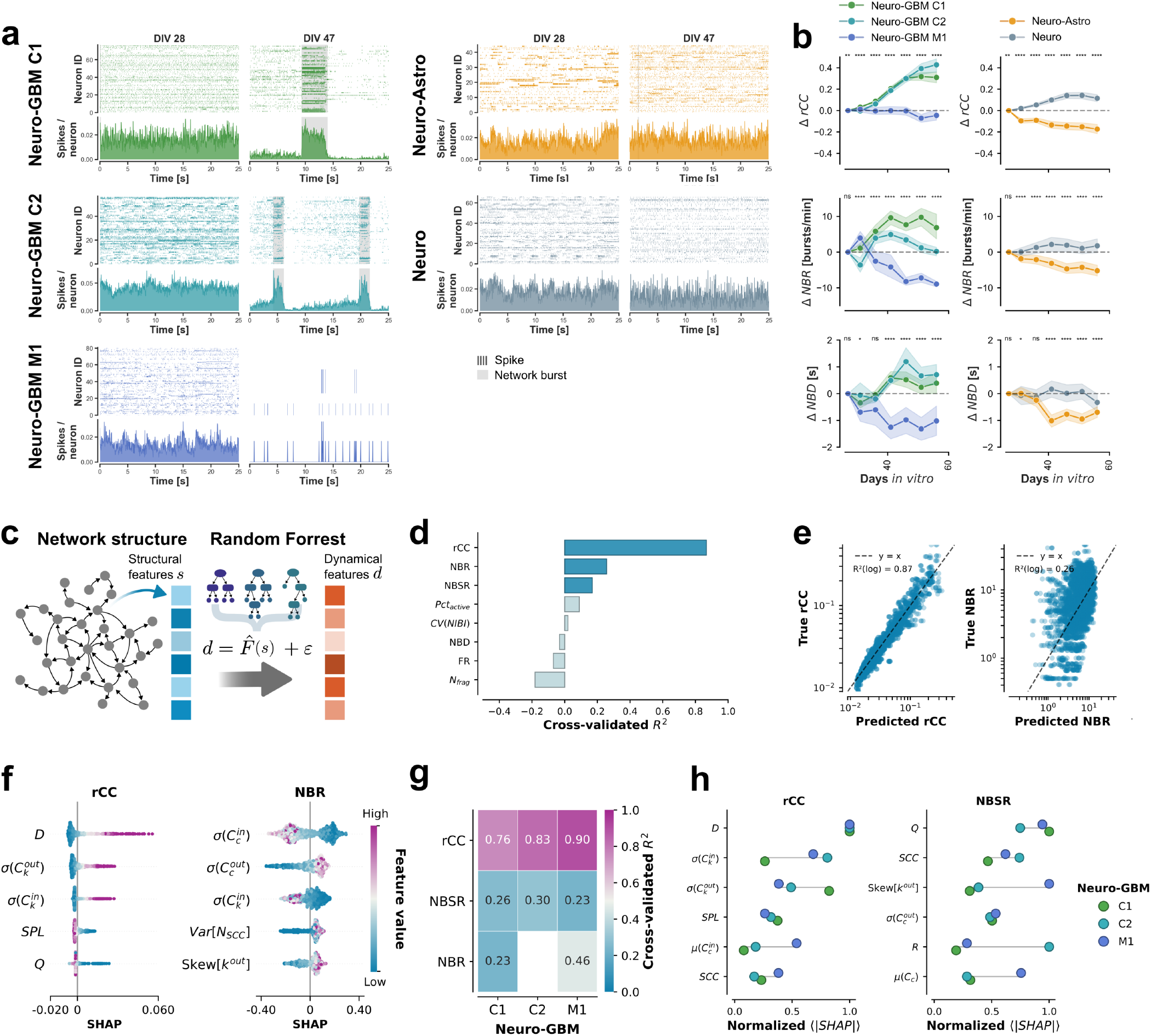
Effective network structure predicts population bursting dynamics. **a** Representative spike rasters (top) and population firing rate (bottom), for the five culture conditions at an early (DIV 28) and a late (DIV 47) timepoint. Classical GBM co-cultures develop pronounced bursts. The mesenchymal type shows near-complete loss of activity, while controls remain comparatively stable. **b** Longitudinal change relative to baseline in pairwise synchrony (Δ*rCC*, top), network burst rate (Δ*NBR*, middle), and network burst duration (Δ*NBD*) over days *in vitro*. Lines and shaded bands show mean ± 95% CI; Data binned in 5 DIVs. Stars indicate significance at each timepoint (Kruskal–Wallis across conditions followed by Dunn’s post-hoc with Benjamini-Hochberg correction, ns, not significant; \**p <* 0.05; \*\**p <* 0.01; \*\*\*\**p <* 0.0001). **c** A random forest regressor maps a fixed set of structural features *s* of the effective network to dynamical features *d*. **d** Cross-validated *R* ^2^ for predicting each dynamical feature from network structure (pooled across cultures). *rCC* is predicted well; *NBR* and within-burst spike rate (*NBSR*) are predicted moderately; the rest of the dynamical features are not predicted. **e** Measured versus predicted values for *rCC* (left) and *NBR* (right); dashed line, identity; *R* ^2^ (log) computed on log-transformed values. **f** SHAP summary (beeswarm) for the *rCC* (left) and *NBR* (right) models for the top 5 features. Each point is one network; horizontal position gives the feature’s signed contribution to the prediction and colour encodes the feature value (low to high). High *rCC* is driven by high network density *D*, large spread in in/out degree centrality 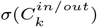, small average shortest path length *SPL*, and low modularity *Q*. High network burst rate is predicted in networks with small variability of in-closeness and in-degree centrality 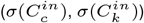 but large spread in out-degree centrality 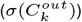 , high variability in strongly connected component size (Var[*N*_*SCC*_ ]), and highly skewed out-degree distribution Skew[*k*^*out*^]. **g** Cross-validated *R*^2^ for predicting rCC, NBSR, and NBR within each patient line (C1, C2, M1); missing entry indicates a model that did not reach predictive performance. **h** Normalized mean absolute SHAP value of the top structural features for the rCC (left) and NBSR (right) models, shown separately for each patient (C1, C2, M1). Predictions rely on a largely consistent feature subset across patients.

To test whether large-scale population dynamics are shaped by connectivity, we trained random forest models to predict individual properties of population bursting from a fixed set of structural features normalized by network size (Fig. 4 c; see “Random forest models” in Methods for training details). These models show that network structure predicts pairwise synchrony (rCC), network burst rate (NBR), and within-network burst spike rate (NBSR), but carries essentially no information about overall firing rate, temporal variability, or fragmentation (Fig. 4 d–e). Cross-validated predictions of rCC closely tracked the measured values (*R*^2^ ≈ 0.87), whereas NBR and NBSR were captured only coarsely (*R*^2^ ≈ 0.26 and *R*^2^ ≈ 0.17, respectively). Synchrony is therefore largely set by network architecture, whereas NBR and NBSR appear to be additionally governed by biophysical mechanisms that structure alone does not capture.

To identify which features drive the well-predicted targets, we decomposed each model’s output into additive per-feature contributions using SHAP (SHapley Additive exPlanations) values (51), which quantify how much each feature pushes a prediction above or below its baseline (Fig. 4 f). rCC is highest in dense networks with short path lengths *SPL*, low modularity *Q*, and high variability in degree centralities 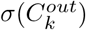 and 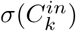 meaning that well-integrated networks with heterogeneous hubs promote synchrony. The NBR follows a different signature: it is elevated when in-closeness 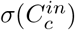 and in-degree centrality 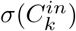 are narrowly distributed but out-closeness centrality 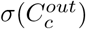 is highly variable. This implicates a broadcasting topology, in which a few neurons with high out-closeness disseminate activity efficiently while their inputs remain homogeneously distributed. These dependencies hold within individual patients (Fig. 4 g), and in all cases the models strongly rely on only a small subset of the available structural features (Fig. 4 h).

This suggests that network burst dynamics are shaped by both network architecture and the biophysical properties of individual neurons, motivating a closer examination of how single-neuron position within the network determines temporal recruitment during bursts.

### Local network properties determine neuronal spatiotemporal recruitment during bursts

A network burst is not a homogeneous event: it has internal temporal structure, with some neurons igniting it and others recruited only later (52). Examining activity at single-neuron resolution during and between network bursts revealed three reproducible firing phenotypes (Fig. 5 a-b and Supplementary Fig. S4 c). To quantify them, we compared each neuron’s firing rate inside network bursts (*FR*_*in*_) with its rate outside (*FR*_*out*_) and combined the two into a network burst-preference index (PI), Eq. 1, which ranges from −1 (firing exclusively outside network bursts) to +1 (firing exclusively within network bursts) (Fig. 5 c). This separated neurons into *bursters*, (*PI* → 1); *tonic* neurons, (*PI* → −1); and *mixed* neurons, which fire comparably in both regimes. The composition of these populations diverged across conditions over development. In classical GBM co-cultures, the fraction of burster neurons rose progressively, whereas it declined in mesenchymal GBM co-cultures as well as in neuron-astrocyte controls, and remained stable in neuron-only controls (Fig. 5 d, top). Bursters also became the earliest-firing neurons in each event: their first-spike latency (FSL) relative to the rest of the population grew increasingly negative in Neuro-GBM C1 and C2 but not in controls (Fig. 5 d, bottom), indicating that in these co-cultures bursters increasingly initiate the bursts.

**Fig. 5.**
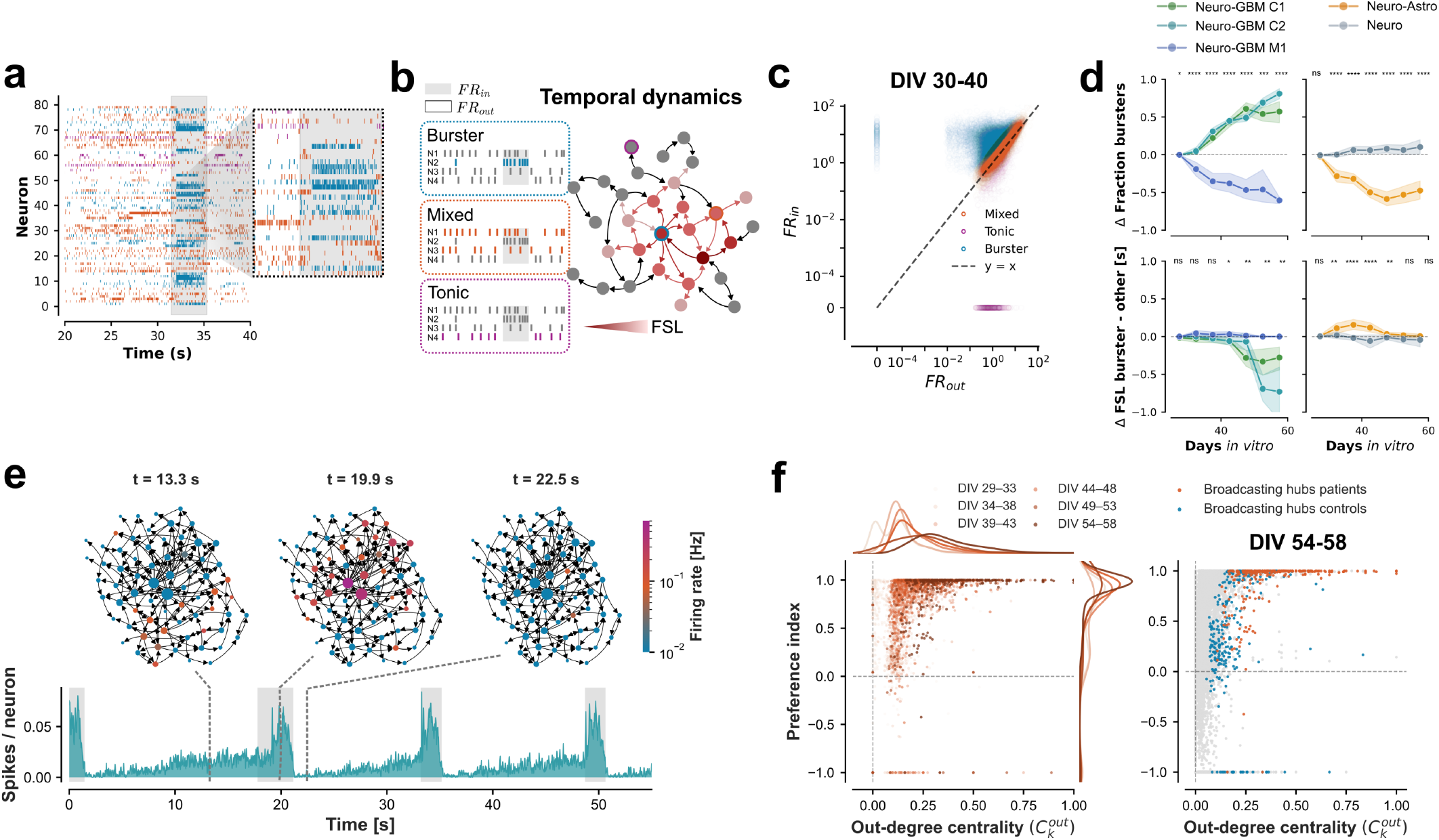
Neurons have distinct temporal roles in population bursts, set by their network position. **a** Representative spike raster with spikes coloured by temporal role. **b** Each neuron’s firing rate inside bursts (*FR*_*in*_) is compared with its rate outside bursts (*FR*_*out*_), yielding bursters, mixed, and tonic neurons. First-spike latency (FSL; red shading) relative to burst onset was calculated for each class, characterising their temporal recruitment during network bursts. **c** *FR*_*in*_ versus *FR*_*out*_ across neurons (DIV 30 to 40), coloured by role; dashed line, identity. **d** Longitudinal change relative to baseline in the fraction of burster neurons (top) and in the first-spike latency (FSL) of bursters relative to other neurons (bottom) over days *in vitro*. Lines and shaded bands show mean ± 95% CI; Data binned in 5 DIVs. Stars indicate, within each panel, a significant difference between conditions at that bin (Kruskal–Wallis across conditions followed by Dunn’s post-hoc, ns, not significant, *p >* 0.05; \**p <* 0.05; \*\**p <* 0.01; \*\*\**p <* 0.001; \*\*\*\**p <* 0.0001). **e** Effective network at three timepoints around a network burst (nodes coloured by firing rate), with the corresponding population rate below (network bursts shaded). Network bursts ignite focally and propagate across the network. **f** Left, burst-preference index versus out-degree centrality 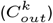 across successive developmental windows (colour, DIV epoch); marginal distributions shown. High-preference neurons progressively accumulate at high out-degree centrality. Right, the same at DIV 54 to 58, highlighting broadcasting hubs in patients (*n* = 266) and controls (*n* = 297): patient hubs are overwhelmingly bursters, control hubs are not.

Finally, we asked whether this temporal leadership has a structural basis. Indeed, central hub neurons sustained high firing rates throughout network bursts, while more peripheral neurons fired predominantly outside of them (Fig. 5 e). Plotting each neuron’s network burst-preference index against its out-degree centrality 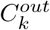 across successive time bins revealed a progressive accumulation of high-preference neurons at high out-degree centrality in patient co-cultures (Fig. 5 f, left). By late stages (DIV 54 to 58), the broadcasting hubs of patient networks predominantly fired during bursts, whereas those of control networks did not develop this tendency (Fig. 5 f, right). Thus, only in GBM co-cultures do the neurons that lead bursts also become the neurons that broadcast most widely. This reveals a GBM-specific convergence of firing behaviour and structural influence.

### Glioblastoma shifts neuronal information processing from unique to redundant integration

Beyond altered network activity such as epileptiform bursting, GBM is associated with progressive cognitive impairment (26, 27), suggesting that tumour-driven network changes carry computational consequences. Having established that GBM reorganises both network topology and the functional roles of individual neurons, we therefore asked whether these changes measurably alter the computational properties of the network. Rather than benchmarking performance on a specific task, which is biased by the choice of task (56) and difficult to implement in *in vitro* networks, we used information dynamics (57, 58) to quantify these properties in a task-independent manner. We focused on how neurons modify information (partial information decomposition, PID (58, 59)) and store it locally (active information storage, AIS (60)).

We applied PID to triplets of neurons, decomposing the total mutual information that two source neurons (*X*_1_, *X*_2_) carry about a target neuron (*Y*) into four non-negative components: unique information from each source (*Unq*(*X*_1_; *Y*), *Unq*(*X*_2_; *Y*)), redundant information shared by both sources (*Red*(*X*_1_, *X*_2_; *Y*)), and synergistic information that only arises from their combination (*Syn*(*X*_1_, *X*_2_; *Y*)) (Fig. 6 a). Total mutual information decreased progressively in classical Neuro-GBM C1 and C2 co-cultures, while remaining stable in controls (Fig. 6 b). Decomposing this change revealed that unique information decreased significantly in Neuro-GBM C1 and C2, while both redundancy and synergy increased symmetrically (Fig. 6 c). Consistent with the observed increase in clustering coefficient, AIS also increased in GBM co-cultures over the co-culture period (Supplementary Fig. S6), in line with previously reported relationships between local clustering and AIS (60).

**Fig. 6.**
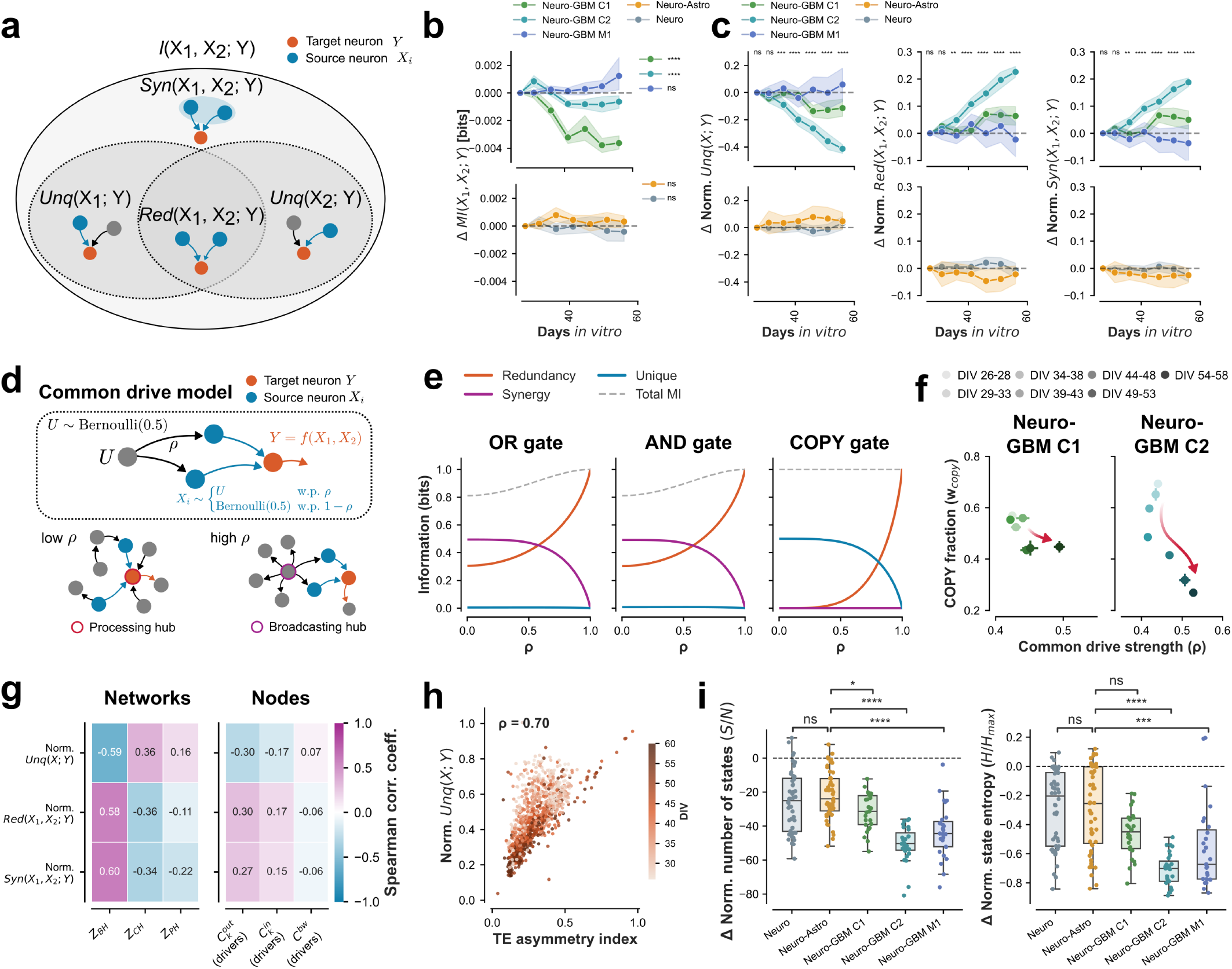
Information-theoretic analysis reveals that glioblastoma invasion restructures neuronal information processing. **a** Partial information decomposition was performed on all identified triplets, decomposing total mutual information into synergistic *Syn*, unique *Unq*, and redundant *Red* components. Venn diagram illustrates the relative distribution of these components. **b** Changes in mutual information *MI* and **c** normalised information contributions relative to baseline recordings, tracked across identified triplets over days *in vitro*. Lines and shaded bands show mean ± 95% CI; Data binned in 5 DIVs. Asterisks indicate significant differences: within panel b, between timepoints within the same condition; within panel c, between conditions at each time bin (Kruskal–Wallis with Dunn’s post-hoc and Benjamini–Hochberg correction, ns, not significant, p > 0.05; *p < 0.05; **p < 0.01; ***p < 0.001; ****p < 0.0001). **d** Schematic of the common drive model used to simulate changes in information contributions. Two source neurons *X*_*i*_ receive input from a shared upstream drive *U* with common drive strength *ρ*. The activity of target neuron *Y* is determined by the source neurons via a gate function *f* . Low *ρ* represents largely independent input streams; high *ρ* represents highly correlated, redundant input. **e** Analytical solutions obtained for different gate types across the range of common drive values *ρ*. **f** Optimal solution space identified for each network, with mean ± SEM shown in bins of 5 DIV. **g** Spearman correlation coefficients between information contributions and network-level hub metrics (broadcasting hub score *Z*_*BH*_ , communication hub score *Z*_*CH*_ , and processing hub score *Z*_*P H*_) and between information contributions and node-level metrics of the source neuron drivers (out-degree centrality 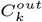, in-degree centrality 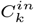, and betweenness centrality 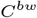. Correlations were considered weak (0.1 ≤ |*ρ*| *<* 0.3), moderate (0.3 ≤ |*ρ*| *<* 0.5), or strong (|*ρ*| ≥ 0.5), following (53). **h** Scatter plot showing the relationship between mean transfer entropy (TE) asymmetry index and normalised unique information. Each point represents one network at one timepoint; only patient-derived networks (Neuro–GBM C1, C2, and M1) are included. Colour encodes DIV. Spearman correlation coefficient *ρ* is shown. **i** Change in the number of identified network states (in 20 ms windows following (54, 55), normalised by total neuron count) and change in state entropy (normalised by the maximum possible entropy given the number of observed states), both computed as the change between DIV 28 and DIV 56/58. Individual networks are shown as scatter points. Asterisks indicate significance (Kruskal–Wallis with Dunn’s post-hoc and Benjamini–Hochberg correction, ns, not significant, p > 0.05; *p < 0.05; **p < 0.01; ***p < 0.001; ****p < 0.0001).

While an increase in redundancy has been previously observed in hyperexcitable or hypersynchronous network states (61), the concurrent rise in synergy implies a more complex underlying mechanism, as correlated activity does not straightforwardly reduce synergistic interactions (62). Identifying the mechanism behind this joint shift would connect the structural reorganisation we observe to its computational consequences, and could potentially inform strategies to preserve information-processing capacity. We hypothesised that the emergence of broadcasting hubs mechanistically drives this shift. In a triplet (*X*_1_, *X*_2_ → *Y*), a broadcasting hub upstream of both sources creates a shared drive that correlates *X*_1_ and *X*_2_, parametrised by the common drive strength *ρ*. When *ρ* = 0, each source carries a private signal to *Y* , giving rise to unique information as in a processing hub integrating independent input streams. As *ρ* increases, sources become correlated, their individual contributions overlap, and information is redistributed from unique toward integrative components. We formalised this as a minimal common drive model in which two source neurons receive input from a shared upstream neuron *U* with coupling strength *ρ*, and the target *Y* computes a logic gate function over its inputs (Fig. 6 d). The PID of any triplet is then fully determined by the gate type of *Y* and *ρ*. We focus on AND, OR, and COPY gates, which are relevant in our experimental setup: COPY gates represent triplets with one dominant source input and an effectively silent second source; AND gates correspond to low-excitability neurons that require convergent input from both sources; and OR gates correspond to high-excitability neurons that fire in response to either source alone. Analytical solutions for OR, AND, and COPY gate computations confirm the expected behaviour: increasing *ρ* redistributes information from unique toward redundant components (Fig. 6 e and Supplementary Fig. S7 a,b). However, increasing *ρ* alone cannot reproduce the observed symmetric increase in both redundancy and synergy. This implies that the observed PID shift reflects both an increase in *ρ* and a change in gate composition, specifically a transition from COPY gate-dominated processing toward OR/AND gate-dominated integration. This could represent a shift from asymmetric connections (COPY) toward correlated integration (OR/AND) that is consistent with GBM-induced hyperexcitability progressively recruiting neurons into a hub-driven activity regime. To quantify this transition, we mapped all (*ρ, w*_*copy*_) pairs consistent with the empirical PID fractions at each time point by minimising the squared residuals (example for the population means at DIV 28 and DIV 58 shown in Fig. S7 c,d). In Neuro-GBM C2, the *w*_*copy*_ decreased from 0.69 to 0.27 over the co-culture period, accompanied by an increase in *ρ* from 0.44 to 0.53 (Fig. 6 f).

Neuro-GBM C1 showed the same trend with smaller magnitude, consistent with its more subtle PID changes.

We then sought empirical support for this mechanistic interpretation. At the network level, broadcasting hub score *Z*_*BH*_ correlated positively with both redundancy and synergy (Spearman *ρ* ≈ 0.6), while processing hubs showed no independent correlation with any PID component (Fig. 6 g). Additionally, there was a moderate correlation (Spearman *ρ* ≈ 0.3) between the out-degree centrality 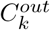 of the neurons driving the source neurons in the triplets and the redundant and unique information contributions. This is consistent with the model and previous literature (63): processing hubs integrate inputs, but what determines the PID structure is the correlation among those inputs, which is set by upstream broadcasting neurons. Additionally, since COPY gates predict strongly asymmetric transfer entropy (one source dominating) and AND/OR gates predict approximately equal contributions from both sources (Fig. S7 e), we used the TE asymmetry index as an empirical proxy for gate composition. The TE asymmetry index is defined as the absolute difference in normalized transfer entropy between the two sources and their target, divided by their sum, ranging from 0 (both sources contribute equally, AND/OR-like) to 1 (one source dominates, COPY-like). The TE asymmetry index decreased progressively over time for Neuro-GBM C1 and C2 cocultures (Fig. S7 f), and correlated strongly with normalised unique information (Spearman *ρ* = 0.70; Fig. 6 h; all PID measures shown in Supplementary Fig. S7 g), providing independent empirical support for the gate composition shift. These shifts in information processing mode had measurable functional consequences. As unique integration gives way to redundant integration, the network loses the diversity of input combinations that individual neurons can independently encode, reducing the number of distinguishable network activity patterns. To quantify this, we mapped population spiking activity onto a sequence of discrete binary states and computed the Shannon entropy of their empirical probability distribution as a measure of the dynamical repertoire (54, 55, 64). Accordingly, both the number of distinct network states visited and the normalised state entropy decreased significantly in Neuro-GBM C1 and C2 co-cultures relative to controls (Fig. 6 i), indicating a progressive collapse of the dynamical repertoire.

## Discussion and Conclusions

While glioblastoma has traditionally been viewed as a focal lesion, mounting evidence reframes it as a network-level disease. Yet, how the tumour reorganises neuronal networks at its periphery, and how this unfolds over time, has remained difficult to study, as longitudinal access to evolving network structure is largely beyond the reach of *in vivo* approaches. Here, we address this with a compartmentalised *in vitro* platform in which iPSC-derived human neurons are co-cultured with primary glioblastoma cells, allowing us to follow network behaviour across the full course of the disease. Because our design forces the connecting axons between compartments to pass in the vicinity of electrodes, we can record from a large fraction of the neurons simultaneously and reconstruct the effective network. This reveals a reproducible mesoscale reorganisation that follows a defined trajectory: an initial shift toward a hyperconnected, hub-dominated architecture, followed by a collapse of community structure that accompanies widespread, non-specific neuronal death. Crucially, these structural changes are not inert. They coincide with a shift towards more synchronisation and sustained network bursting. At the computational level, the network shifts from unique toward redundant and synergistic information processing, and its accessible dynamical state space collapses.

The progressive network reorganisation is captured by simple network-growth rules, namely preferential attachment together with triad formation. A plausible biological substrate for this is activity-dependent synaptogenesis (65): GBM-induced hyperexcitability could selectively strengthen connections onto already-active neurons, so that the most active neuronal units progressively become the most connected. The resulting hub-dominated topology may favour tumour progression: by sustaining network-wide activity, it could promote release of neuronal signalling factors driving tumour growth, such as brain-derived neurotrophic factor and neuroligin-3 (66, 67), and activate neuron-to-glioma synapses (9, 68). Furthermore, this coupling between activity and connectivity may be self-reinforcing: In diseased networks, broadcasting hubs become preferentially active during bursts; if these recruited neurons strengthen their connections via activity-dependent synaptogenesis, structure and dynamics would mutually reinforce. Such positive feedback is normally kept in check by homeostatic mechanism that scale synaptic strength to maintain a stable activity set point (69), but these regulatory processes may be disrupted in disease (70), allowing hub dominance to grow uncompensated.

Hub-dominated, low-modularity networks have a lower barrier to large-scale percolation and contagion (71–73), and in epileptic brain networks, synchronisation of highly connected hubs is a central driver of seizure generation and propagation (22). Computational modelling confirms that even a small number of such hubs is sufficient to render a network seizure-prone without any change in overall connection number (74). The population bursts that dominate our late-stage co-cultures are hypersynchronous and become more frequent and sustained than in controls, hallmarks of the hypersynchronous discharges that characterise seizures *in vivo* (75). The identified structural changes therefore offer a networklevel mechanistic account of GBM-associated epilepsy that complements existing molecular explanations (76, 77), and yield a concrete, testable prediction: if replicated *in vivo*, hub dominance and low modularity measured at diagnosis should predict seizure propensity, providing candidate network markers for the extent of glioblastoma-neuron functional integration.

The accompanying changes in network computation point to a parallel account of GBM-related cognitive decline. A reduced dynamical repertoire, defined as the collapse of diverse activity patterns into a restricted set of recurring states, is a signature of disrupted neuronal computation across pathological and low-information brain states (64, 78). Our information-theoretic results make this concrete: the shift from unique toward redundant and synergistic processing reflects a loss of specificity in neural coding, while the contraction of the dynamical state space and its entropy reflects a loss of flexibility. This computational restructuring is mechanistically driven by a transition from asymmetric COPY gate-dominated processing toward integrative AND/OR gate processing as hub-mediated common drive increases. Although AND and OR gates differ in excitability threshold, both are plausibly promoted by GBM-induced changes: OR gates by glutamatergic hyperexcitability (10), and AND gates by the convergent hub-driven input streams that emerge as broadcasting hubs dominate. Crucially, because this collapse follows from network topology rather than from local cellular changes alone, it offers an explanation for why cognitive symptoms can persist after resection. One possibility is to exploit network control theory (79) to identify patterns of targeted stimulation that restore modularity or curb hub dominance. Delivered through approaches such as responsive neurostimulation (80, 81), this could potentially preserve information-processing capacity.

Several limitations define clear next steps. Our cultures contained only excitatory neurons. Incorporating inhibitory and glial populations would yield a more faithful model, especially since recent studies have pointed to the selective loss and dysfunction of peritumoral GABAergic interneurons as a key driver of GBM-associated hyperexcitability (28, 82). The two GBM subtypes also showed pronounced differences in their temporal dynamics: The mesenchymal subtype showed faster neuronal loss, consistent with its reported lower median survival *in vivo* (40), and an absence of the hub-dominated network restructuring seen in classical subtypes. This subtype specificity warrants dedicated investigation with larger cohorts. Our analyses rest on short (two-minute) recordings, which may miss slower or transient shifts in activity; the continuous, long-term recordings now enabled by emerging recording technologies (35) would capture the full timescale of network reorganisation and reveal dynamics inaccessible to brief snapshots. Most importantly, the platform establishes correlations between structural and functional change but not their direction: causal interventions, such as optogenetic silencing of hub neurons, will be needed to confirm that topological reorganisation drives the dynamical and computational changes we observe. Beyond establishing causality, the same interventional access makes the platform a testbed for pharmacological approaches designed to reduce network-wide hyperexcitability. Finally, relating these signatures to neuroimaging-derived connectivity or testing whether the topological signatures we identify track cognitive decline in patients is a promising avenue to extend these findings to clinical relevance.

By establishing an *in vitro* platform for the longitudinal study of tumour–neuron interaction, this work opens a route to insights into GBM progression and offers a tool of immediate practical value for drug screening and patient-specific modelling. Our central finding is that GBM drives a progressive mesoscale reorganisation of neuronal networks that systematically degrades their capacity to process information, tightly coupling the structural and functional consequences of tumour infiltration. This principle need not be specific to GBM; it could apply to any pathology that reshapes network topology in the same way. Indeed, comparable topological signatures recur across epilepsy, Alzheimer’s disease, and other conditions marked by hyperexcitability (20, 21), suggesting that such measures may serve as disease-agnostic markers of circuit dysfunction. More broadly, the convergence of network neuroscience with engineered, patterned *in vitro* networks marks a promising new frontier, with which structural rules governing neural computation, and their breakdown in disease, can be observed, perturbed, and understood.

## ACKNOWLEDGEMENTS

GA and VV were supported by ETH Zurich and Swiss National Science Foundation (Project Nr. 10001282). KA acknowledges support from Strategic Basic Research Foundation Flanders (FWO) fellowship (1S03424N) and Swiss Government Excellence Scholarship (2024.0119). CMT was supported by ETH Postdoctoral Fellowship. GH acknowledges support from Swiss National Science Foundation Professorial Fellowship (PP00P3_176974); Swiss National Science; Foundation Project grant (3200-0-239894); the ProPatient Forschungsstiftung, University Hospital Basel (Annemarie Karrasch Award 2019); Swiss Cancer Research Grant (KFS-4382-02-2018); the Department of Surgery, University Hospital Basel. The authors further thank LBB members for fruitful discussions.

## Data and material availability

Data will be made publicly available with the final publication. Code is available at https://gitlab.ethz.ch/amosg/gbm_neuro_info.git.

## CRediT author contributions GA

conceptualisation, methodology, supervision, software, validation, formal analysis, investigation, visualisation, and writing – original draft, writing – review & editing. LJ: methodology, validation, investigation, and writing – review & editing. KA: methodology, investigation, and writing – review & editing. AG: resources, methodology, and writing – review & editing. GH: resources, writing – review & editing. JV: conceptualisation, resources, project administration, funding acquisition, and writing – review & editing. CMT: conceptualisation, methodology, supervision, validation, investigation, and writing – review & editing. VV: conceptualisation, methodology, supervision, software, validation, formal analysis, investigation, visualisation and writing – original draft, writing – review & editing.

## Methods

### Experimental methods

#### Design of PDMS microstructures

PDMS microstructures were designed in AutoCAD (Autodesk) and fabricated via soft lithography by WunderliChips GmbH (Zurich, Switzerland). Each microstructure accommodates nine spatially separated networks in parallel (Figure 2 b). Within each network, three seeding wells are connected by two arrays of ten microchannels, each measuring 10 *µ*m in width, 4 *µ*m in height, and 500 *µ*m in length. The seeding wells are ≈ 250 *µ*m in both height and diameter, providing sufficient space to contain and immobilize spheroids, which are smaller in size. The microchannel entries include 1.5 *µ*m-wide constrictions (“Somatic constrictions”, Figure 1 a) designed to reduce glioblastoma migration toward the central well. The channel cross-sectional dimensions (10 × 4 *µ*m) also increase electrical resistance, improving signal detection on underlying electrodes.

#### Preparation of PDMS microstructures

PDMS microstructures were cut from the wafer, immersed in isopropanol for at least 30 min and rinsed five times with sterile ultrapure water (Millipore Milli-Q System, 18.2 MΩ·cm). Following aspiration of excess liquid, the microstructures were left to dry at room temperature for a minimum of one hour. The dried microstructures were placed either on glass-bottom dishes (35 mm, KIT-3522 T, WillCo Wells) for imaging or on high-density CMOS microelectrode arrays (MaxOne, MaxWell Biosystems AG, Switzerland) for electrophysiological recordings. Prior to use, both substrate types underwent plasma cleaning for 2 min (18 W, PDC-32G, Harrick Plasma), were coated with 0.1 mg/mL poly-D-lysine (PDL, Sigma-Aldrich, Cat. No. P6407) in phosphate-buffered saline (PBS, Gibco, Cat. No. 10010015), rinsed three times with ultrapure water, and blow-dried with nitrogen. For MEA recordings, the microstructures were carefully aligned over the 3.85 × 2.10 mm sensing area using tweezers. To promote adhesion of the PDMS to the glass and CMOS chips, the dishes were briefly desiccated for 2 min, before being placed in an oven at 37 °C for at least 30 min. 1 mL of pre-warmed PBS was added to the dish and the assembly was desiccated to remove any air trapped within the microchannels.

#### Cell cultures

##### Neuron culture

Cryogenized aliquots of human induced pluripotent stem cell (iPSC)-derived NGN2 Neurons (hDFa90 /1.2iNgn2 p8+35; Novartis) were kindly provided by Novartis and stored in liquid nitrogen until use. NGN2 neurons were thawed and aggregated into spheroids at a density of 200 cells per spheroid using AggreWell^TM^ 400 microwell plates (StemCell Technologies, 34415) according to the protocol of the manufacturer. During spheroid formation, cells were maintained in combined Neuron-Glioblastoma (CNG) medium prepared from Neurobasal plus (Thermo Fischer Scientific, A3582901) medium with glutamax (100 X, Gibco, 35050061), sodium pyruvate (100 X, Gibco, 11360070), antibiotic-antimycotic (100 mM, Gibco, 15240096), N2 supplement (100 X, Gibco, 17502048), B27 (50 X, Gibco, 17504044), BDNF (10 ng/mL, PeproTech, 450-02) and GNDF (10 ng/mL, Preprotech, 450-51), supplemented with doxycycline (1 *µ*L/mL, Fisher Scientific, NC0424034), laminin (5 *µ*g/mL, Sigma-Aldrich, L2020-1MG), and rho-kinase inhibitor (10 *µ*M, Sigma-Aldrich, Y27632, 688000). Approximately 24 hrs after thawing, neuronal spheroids were seeded in the mounted PDMS microstructures. After seeding, cultures were maintained in CNG medium supplemented with doxycycline (1 *µ*l/mL) and laminin (5 *µ*g/mL) for the first week *in vitro* (DIV 0–DIV 7). From DIV 7 to DIV 10, the medium was supplemented only with laminin, and from DIV 10 onward, cultures were maintained in CNG medium alone.

##### Glioblastoma culture

Primary glioblastoma cells were derived from tumour tissue resected from patients undergoing surgery at the University Hospital Basel, Switzerland (see “Ethics statement”). Using DNA methylation–based classification (37), as applied in neuropathological diagnosis (38, 39), the patient-derived tumour cells were assigned to the classical and mesenchymal subtypes.

The primary patient-derived GBM cells were thawed and kept in 25 mL flask for minimum one week before use. The cells were maintained in Neurosphere (NS) medium, prepared by adding glutamax (100 X, Gibco, 35050061), sodium pyruvate (100 X, Gibco, 11360070), non-essential amino acids (100 X, Gibco, 11140050), sodium bicarbonate (7.5%, Gibco, 25080094), HEPES buffer 30mM (1M in *H*_2_0, C-40020, Sigma Aldrich), antibiotic-antimycotic (100X, Sigma Aldrich, A5955), EGF (0423AF05, Sigma-Aldrich, SRP3027-500UG), FGF (154 a.a., Peprotech, 100-18B-1MG), heparin (1000 X, StemCell Technologies, 07980) and B27 w/o vitamin A (50 X Gibco, 12587001) to 46.75 mL of neurobasal medium (Gibco, 21103049) and 46.75 mL DMEM /F12 (Gibco, 11320033). Prior to spheroid preparation, primary glioblastoma samples in culture were collected and resuspended in 5 mL of TrypLE (Cat.No: 12604021, Gibco, ThermoFisher) and 50 *µ*L of DNAse (Cat. No: EN0521, ThermoFisher) and incubated for 15 minutes at 37°C. Manual dissociation was performed to obtain a single cell solution. The cells where then aggregated into spheroids at a density of 100 cells per spheroid using AggreWell^TM^ 400 microwell plates.

##### Astrocyte culture

Astrocyte progenitors were generated by inducible overexpression of the transcription factor SOX9 in the Sigma iPSC line Sigma-iSOX9-iPSC0028, following the protocol described by Neyrinck et al. (83). iPSCs were maintained in E8 Flex medium for 3-5 days prior to dissociation using accutase and replating at a density of 100,000 cells/cm^2^ on Matrigel-coated 6-well plates. On day 1 postplating, cells were cultured in neural maintenance medium (NMM), prepared with 1:1 mixture of Neurobasal medium (Gibco) and DMEM /F12 supplemented with 0.5 X gluta-max, 50 *µ*/mL Penicillin-Streptomycin, 0.5 X B27 (Gibco), 0.5X N2 (Gibco), 0.5 X MEM-NEAA (Gibco), 0.5 X sodium pyruvate, 0.025 % human insulin (Sigma) and 50 µM 2-mercaptoethanol (Gibco). NMM was further supplemented with 1 µM LDN193189 and 10 µM SB431542. The media was changed daily.

After 12 days of induction, cells were dissociated using accutase and replated at 65,000 cells/cm^2^ on Matrigel-coated plates in NMM supplemented with 1 X RevitaCell and 20 ng/mL basic FGF. The following day, medium was switched to astrocyte maturation medium (AMM), composed of 1:1 mixture of Neurobasal medium and DMEM /F12 supplemented with 1 X N2, 1 X sodium pyruvate, 1 X glutamax, 0.5 mM N-acetyl-cysteine (Sigma), 0.1 mM dbcAMP (Sigma), 10 ng/mL CNTF (Peprotech), 10 ng/mL BMP4 (Peprotech) and 5 ng/mL HB-EGF (Peprotech). AMM was further supplemented with 3 *µ*g/mL doxycycline to induce SOX9 expression. Medium was replaced every other day. Cells not intended for further use were cryopreserved in AMM supplemented with 20 % dimethyl sulfoxide (DMSO) at day 18.

On day 21, the full medium was replaced with CNG medium to allow the cells time to acclimatise. The following day, cells were detached using accutase and aggregated into spheroids at a density of 100 cells per spheroid using AggreWell^TM^ 400 microwell plates. The differentiated astrocyte progenitors were characterised by immunostaining with GFAP, S100*β*, and SOX9 antibodies after 6 days of culture in CNG medium, to confirm that the culture medium did not affect differentiation (Fig. S1 e–f).

##### Cell seeding

1 h prior to cell seeding, the PBS contained in the substrates was replaced with 1 mL of CNG media and placed in the incubator. Two neighbouring wells in each of the nine networks were manually seeded with one neuronal spheroid per well using a 10 *µ*L pipette under a bench-top microscope. Following a 28-day maturation period, the CNG media was replaced by CNG+ media. CNG+ was used as a coculturing media to support glioblastoma growth and was prepared by supplementing CNG media with EGF (0.2 µL/mL), FGF (0.8 µL/mL), and Heparin 1000 X (1 *µ*L/mL). Then, a single glioblastoma spheroid respectively a single astrocyte spheroid for positive controls was introduced into each of the remaining empty wells.

##### Culture maintenance

To support long-term culture, medium was refreshed every 2–3 days by replacing half of the volume with fresh medium.

#### Imaging

##### Cell tracking

To visualize interactions between glioblastoma cells and mature neuronal networks, glioblastoma spheroids were labelled with CellTracker Orange CMRA (1 *µ*l/mL, ThermoFisher, C34551), which was added directly to the spheroid culture medium in the AggreWell^TM^ plates without disturbing the spheroids prior to seeding. Neuronal cultures were stained in parallel with CellTracker Green CMFDA (1 *µ*l/mL, ThermoFisher, C7025). Both cell types were incubated with the dyes for 40 min and subsequently rinsed by two complete medium exchanges. After seeding glioblastoma spheroids into the networks, cultures were imaged on an Olympus Fluoview 3000 confocal microscope using a 10X objective (NA 0.3), with images acquired every 30 min.

##### Immunostaining

Cultures were fixed in 4 % paraformaldehyde (Sigma-Aldrich, 1004960700) for 15 min at room temperature, followed by three 10 min washes in PBS. Cells were then permeabilized with 0.1 % Triton X-100 (Sigma-Aldrich, 9002931) for 6–8 min at room temperature. To block non-specific binding, samples were incubated in blocking buffer (5 % goat serum in 1X PBS, ThermoFisher, 50197Z) with six buffer exchanges over a period of at least ≥ 6 h. Primary antibody incubation was performed overnight at room temperature. The following day, samples were washed six times with PBS over ≥ 6 h and incubated with secondary antibodies overnight at room temperature. Afterward, two 10 min PBS washes were performed, followed by nuclear staining with Hoechst 33342 (1 *µ*l/mL, Invitrogen, H3570) for 40 min. Finally, cultures were washed twice in PBS prior to imaging. The used antibodies are listed in Supplementary Table 5. Images were acquired using an Olympus Fluoview 3000 confocal microscope with 10X, 20X (NA 0.5) and 60X (NA 1.3) objectives. CLSM images were processed using custom macros in the open-source software Fiji (ImageJ2, Version: 2.14.0/1.54f). Synapse puncta quantification was done using SynBot (84). Brightness and contrast were adjusted to improve image clarity for illustrative purposes.

#### Electrophysiology

##### Recordings

**\**Networks were recorded in an incubator at 5 % CO_2_ and 35 °C, corresponding to a recording temperature of 36-37 °C in the media. The cultures were allowed to equilibrate for ≥ 3 min before recording to stabilize activity that could have been affected by handling or environmental fluctuations. Electrodes not covered by the microstructure were excluded based on impedance maps, and ≈ 600 electrodes per network were selected using custom software as previously reported (34). Spontaneous activity was acquired at 20 kHz for 2 min and saved as HDF5 files through a custom-built interface based on the manufacturer’s Python API (MaxWell Biosystems). Baseline activity was measured on DIV 26–28, followed by recordings three times per week after glioblastoma or astrocyte addition.

##### Spike sorting

HDF5 files were processed with a custom Python pipeline. Spikes were detected and sorted using SpikeInterface (85) with SpyKING CIRCUS v2 (86). Thresholds for spike detection were set at five times the signal standard deviation (negative peaks) with a 3 ms minimum interspike interval. Units with SNR *<*5 or firing rate *<*0.1 Hz were excluded. Examples for identified units are shown in Supplementary Figure S4.

##### Functional activity metrics

Per-neuron and network-level functional features are listed with their abbreviations in Supplementary Table 3; standard metrics (firing rate, inter-spike intervals, etc.) were computed using conventional definitions, and we detail below only the quantities requiring specific operational choices. To quantify each neuron’s firing relative to the network burst, we computed the mean first-spike latency with respect to network-burst onset (*FSL*) and a burst-preference index (*PI*),

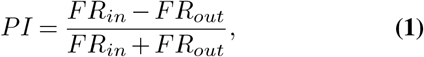

where *FR*_*in*_ and *FR*_*out*_ are the firing rates inside and outside network bursts.

Network bursts were identified from the population firing rate, obtained by binning spikes in 25 ms windows and smoothing with a Gaussian kernel. Network bursts were defined as activity exceeding seven standard deviations above baseline for ≥ 50 ms, involving ≥ 30 % of neurons. Baseline was estimated as the median of the lower 50 % of the smoothed firing rate. Network bursts ended when activity fell below one standard deviation above baseline for ≥ 50 ms. A minimum threshold of 6 Hz was applied to reduce false positives in low-activity cultures. The instantaneous correlation (rCC) was calculated by binning the data in 5 ms windows and averaging the pairwise Pearson’s correlation coefficients between all combinations of *N* binned spike trains using elephant (87).

### Information-theoretic analyses

Spike trains from spike-sorted units were converted into binary spike trains at 1 ms temporal resolution, yielding a binary matrix of dimensions *S* × *T*_bins_, where *S* is the number of units identified by spike sorting, *T*_bins_ is the total number of timesteps in the binned spike trace, and entries of 1 indicate a spike occurrence. All subsequent information-theoretic analyses were performed on these binarised representations using the IDTxl toolbox (88) with the Java Information Dynamics Toolkit (JIDT) (89) as the underlying estimator backend.

All measures reported below are built on (conditional) mutual information. The mutual information between two discrete variables *A* and *B* quantifies the reduction in uncertainty about one given knowledge of the other,

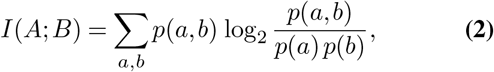

measured in bits, and is zero if and only if *A* and *B* are independent. The conditional mutual information *I*(*A*; *B* | *C*) is the analogous quantity evaluated within the distribution conditioned on a third variable *C*, capturing the dependence between *A* and *B* that remains after accounting for *C*.

#### Inference of effective connectivity using multivariate transfer entropy

To infer connectivity between units, we represented each spike-sorted unit as a node in a weighted directed graph *G* = (*N, ℰ*) and quantified the information flow between units using transfer entropy. A directed edge (*i, j*) ∈ ℰ denotes a significant transfer of information from the source unit *i* to the target unit *j*, with weight *w*_*ij*_ given by the normalised mTE from *i* to *j*. For an ordered pair of units with binarised spike trains *X* and *Y* , a directed edge *X* → *Y* denotes a statistically significant transfer of information from the past of the source unit *X* to the present of the target unit *Y* ; the edge weight encodes the strength of this transfer. Bivariate transfer entropy (biTE) at delay *u* is the information the source’s past provides about the target’s present, conditioned on the target’s own past:

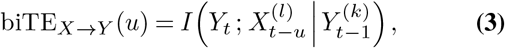

where 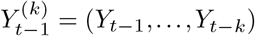 is the *k*-dimensional target past and 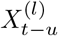 the *l*-dimensional source past at delay *u*. Multivariate TE additionally conditions on the past states **Z**^−^ of all other retained sources of *Y* , isolating the information *X* contributes beyond that already provided by the other inputs:

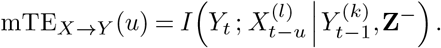

Directed effective connectivity was inferred via the two-stage transfer entropy procedure described in detail in our previous work (90). In the first stage, biTE was computed between all ordered pairs of units using the discrete JIDT estimator, with a target embedding of *k* = 5 bins (5 ms), a source embedding of *l* = 1 bin (1 ms), and delays swept over 1–10 ms retaining the value and the delay at which maximum was retained. Statistical significance was assessed by a permutation test (*n* = 200), and *p*-values were corrected for multiple comparisons using the Benjamini–Hochberg false discovery rate (FDR) procedure. Source–target pairs with FDR-corrected *p* ≤ 0.05 were retained as candidate connections, and the delay maximising significant biTE was recorded for each pair. In the second stage, mTE was computed for each target unit using the `MultivariateTE` class with the same discrete estimator and *k* = 10 bins (10 ms), restricting candidate sources to those identified in the first stage at their optimal biTE delay; final conditioning sets were selected by IDTxl’s greedy iterative procedure, with significance assessed against 200 permutation surrogates and FDR correction (*p* ≤ 0.05). Retained TE values were normalised by the Shannon entropy of the target unit, to control for firing-rate variability (91). Inferred connections were represented as directed, weighted graphs using NetworkX (92).

##### Active Information Storage

Active Information Storage (AIS) quantifies the degree to which a unit’s current activity is predicted by its own past, and was computed for each unit from the binarised spike trains using the `ActiveInformationStorage` class in IDTxl with a discrete CMI estimator (`JidtDiscreteCMI`). For a unit *Y* , AIS is the mutual information between its past state and its current state (93):

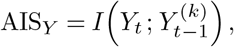

where *Y*_*t*_ is the current state and 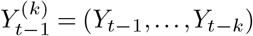 is the *k*-dimensional past, consistent with the embedding notation used above. The embedding dimension *k* was optimised automatically over lags up to a maximum of 10 ms. Statistical significance was assessed using 200 permutations for both maximum and minimum statistic tests. To control for the confounding effect of variable firing rates across units, AIS values were normalised by each unit’s Shannon entropy-yielding a normalised value in the range [0, 1].

#### Partial Information Decomposition

Partial Information Decomposition (PID) was computed for all neuron triplets *X*_1_, *X*_2_ *Y* in which both sources *X*_1_, *X*_2_ were identified as significant mTE sources of the same target *Y* . For each such triplet, bivariate PID was estimated using the `BivariatePID` class in IDTxl with the Sydney PID estimator (94), decomposing the joint mutual information *I*(*X*_1_, *X*_2_; *Y*) into four non-negative components:

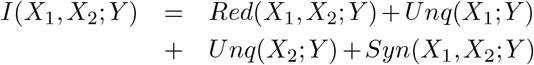

where *Red*(*X*_1_, *X*_2_; *Y*) is the redundant information carried by both *X*_1_ and *X*_2_ individually, *Unq*(*X*_*i*_; *Y*) is the unique information carried only by *X*_*i*_, and *Syn*(*X*_1_, *X*_2_; *Y*) is the synergistic information available only from the combination of *X*_1_ and *X*_2_. Source lags were fixed to the values identified during mTE inference. Each PID component was subsequently normalised by the joint mutual information of the triplet to yield fractional contributions summing to unity, enabling comparison across triplets and conditions with differing firing statistics.

### Validation of effective connectivity inference

To assess the performance of mTE-based network inference relative to simpler methods, cross-correlation (CC), biTE, and mTE were applied to synthetic spike train data with known ground-truth connectivity, and their precision, recall, and false-positive rates were compared as a function of synaptic strength.

#### Ground truth models

We simulated recurrent networks of *N* = 60 excitatory adaptive exponential leaky integrate-and-fire (AdEx) neurons (44) coupled through conductance-based synapses implemented in Brian2 (95) and integrated with forward Euler (Δ*t* = 0.1 ms). Neuron parameters followed the AdEx regular-spiking set (44), with a 2 ms refractory period. To benchmark mTE inference across topologies, we generated three classes of directed ground-truth graphs (*n* = 20 per class): *balanced* graphs, drawn as Erdős–Rényi networks with edge probability *p* ∼ *U* [0.01, 0.05]; and *broadcasting-hub* and *processing-hub* graphs, in which three randomly chosen nodes of an Erdős–Rényi baseline (*p* = 0.025) received additional outgoing or incoming edges to *k* ∼ *U* {5,…, 15} randomly sampled partners. Each edge *i* → *j* carried an integer conduction delay *d*_*ij*_ ∼ *U* {1,…, 10} ms and a structural weight *c*_*ij*_ ∼ *U* [0, 1], giving synaptic conductance *w*_*syn,ij*_ = *wc*_*ij*_. Synapses were excitatory (*E*_exc_ = mV) with a bi-exponential, AMPA-like conductance (rise ms, decay 2 ms); a fraction *f*_dep_ were depressing and the remainder facilitating (96).

Every neuron received constant bias current that was set to a fraction #x223C; *U* [0.7, 1] of the analytically computed AdEx rheobase, and an independent excitatory shot-noise background (*K* = 50 Poisson trains of rate ν, each event injecting *Q* = 0.1 nA and decaying with *τ*_poi_ = 5 ms). For each graph we drew 25 parameter sets, sampling independently and uniformly the mean synaptic conductance *w* ∈ [5, 200] nS, and the Poisson rate ν ∈ [1, 10] Hz. Each network was simulated for 125 s; we discarded the first 5 s as transient and retained recordings with mean firing rate in [1, 20] Hz.

Networks inferred using mTE were compared against two alternative approaches:

- **Cross-correlation** Pairwise CC histograms were computed for all ordered unit pairs using the Elephant library (87), with CC histogram windows of ± 10 ms. For each pair, the peak normalised CC coefficient was extracted separately for positive lags and negative lags. To establish a significance threshold, a null distribution of peak CC values was constructed for each pair by generating n=200 surrogate spike trains with uniformly randomised spike times but identical spike counts. The empirical p-value was computed as the proportion of surrogate peak CC values exceeding the observed value, and connections were declared significant at *α* ≤ 0.05.
- **Bivariate transfer entropy** In addition to its role as the first-stage filter for mTE, biTE was analysed as a stand-alone measure of pairwise directed coupling, computed and thresholded as described above. Unlike mTE, biTE does not condition on other simultaneous inputs and therefore cannot disambiguate direct connections from shared drive or cascade effects.

#### Performance metrics

For each method and each value of synaptic strength *I*_*syn*_, inferred edges were compared against the ground-truth adjacency matrix to compute precision (positive predictive value), recall (sensitivity), and false-positive rate (FPR). Recall was also evaluated as a function of the ground-truth synaptic conductance *w*_*syn,ij*_ of each connection to characterise each method’s sensitivity to weak versus strong connections.

### Network analysis

All graph-theoretic analyses were performed using NetworkX (92). A full list of computed metrics and their abbreviations is provided in Table 4.

#### Neuron-level metrics

For each node, the following metrics were computed. In-degree centrality 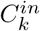 and out-degree centrality 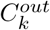 were calculated as the fraction of all possible incoming and outgoing connections that a node participates in, respectively, and thus normalised by *N* −1. Node betweenness centrality 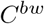 was computed as the fraction of all shortest paths in the network that pass through a given node, normalised over all pairs (97). In-closeness centrality 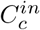 and out-closeness centrality 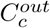 measured the mean inverse shortest path length from all other nodes to a given node, and from a given node to all others, respectively, normalised by *N* − 1. The local clustering coefficient *C*_*C*_ was calculated as the fraction of a node’s neighbours that are also connected to each other, using the directed formulation (98, 99).

#### Network-level metrics

Density was computed as *D* = *E/*(*N* (*N* − 1)), the fraction of realised among all possible directed edges. Clustering was summarised by the mean *µ*_*C*_ and standard deviation *σ*_*C*_ of the local clustering coefficient across nodes. Global transitivity *T* , the ratio of closed triangles to all connected triplets, and reciprocity *R*, the fraction of edges with a reciprocal counterpart, were also computed. The distribution of degree centrality across nodes was described by its mean, standard deviation, skewness, and kurtosis. Degree assortativity was quantified by the directed assortativity coefficients *r*^in-in^, *r*^in-out^, *r*^out-in^ and *r*^out-out^ (100), which measure the tendency of nodes to connect to others of similar degree, computed as the Pearson correlation between the relevant degrees across all directed edges.

Path-based properties were computed on the directed graph over all reachable node pairs. We recorded the mean shortest path length *SPL* and its standard deviation, as well as the graph diameter (the maximum shortest path length). Global efficiency *Eff* was computed as the mean of the inverse shortest path lengths over all ordered pairs. Community structure was identified using the Louvain algorithm (101) on the undirected projection of each network, with modularity *Q* computed over the resulting partition; the number of communities *M* and the fraction of neurons in the largest community *f*_*LC*_ were retained as summary statistics. Strongly connected components (SCCs) were identified on the directed graph, and the number of SCCs, the variance in SCC size, and the fraction of nodes in the largest SCC *f*_*SCC*_ were retained as measures of global directed reachability.

#### Hub neuron detection

Candidate hub neurons were defined as nodes whose centrality exceeded the 95th percentile of the network’s degree centrality distributions. Three functional hub types were distinguished based on the centrality measure used for classification: broadcasting hubs *Z*_*BH*_ , defined by high out-degree centrality and corresponding to neurons that drive many downstream targets; processing hubs *Z*_*P H*_ , defined by high in-degree centrality and reflecting neurons that integrate inputs from many sources; and communication hubs *Z*_*CH*_ , defined by high betweenness centrality and reflecting neurons that lie on many shortest paths between other node pairs (102). To assess whether the degree of each hub neuron candidate exceeded chance expectation, a null distribution was constructed by generating *n* = 500 Erdős–Rényi random graphs (103) matched to the observed network in number of nodes *N* and edge probability *p* = *E/*(*N* (*N* −1)). For each null graph, the mean centrality of hub-classified nodes was computed for each hub type. The observed mean hub strength was then compared against this null distribution to yield a *z*score and two-tailed empirical *p*-value for each hub type at the node level, asking whether this individual node’s centrality is stronger than expected given its null distribution.

### Separability analysis

To assess how distinguishable each group was from controls across development, we trained binary logistic-regression classifiers within 7-day developmental bins (DIV 28–59). Within each bin we defined a negative class (e.g. pooled controls) and a positive class (e.g. a single patient line) and used the structural features as predictors. Each classifier was an *L*_2_-regularized logistic regression (*C* = 1) with balanced class weights, fit on *z*-scored features. All remaining hyperparameters were left at their SCIKIT-LEARN (v1.7.1) defaults. Separability was evaluated by leave-one-chip-out cross-validation, holding out all recordings from one chip per fold and collecting out-of-fold predicted probabilities; folds with a single class in the training set were skipped. Discriminability was summarized as the area under the ROC curve (AUC) computed over the pooled out-of-fold predictions, where AUC = 0.5 denotes chance.

### Generative Model

#### Empirical data

To investigate the wiring rules underlying network reorganisation, we fitted generative models of directed graph evolution to empirically observed connectivity changes. For each patient, networks recorded at an early time point served as baseline graphs *G*_0_, and networks recorded at a later time point served as target graphs *G*_emp_. The early stage spanned DIV 26–28 (baseline graphs) to DIV 41 (target graphs), and the late stage spanned DIV 41 (baseline graphs) to DIV 58 (target graphs).

#### Goodness-of-fit: energy function

To quantify how well a synthetic graph *G*_syn_ reproduced the observed target graph *G*_emp_, we defined an energy function based on the Kolmogorov–Smirnov (KS) statistic (49). Four topological properties were compared between synthetic and observed graphs: clustering coefficient (*C*_*C*_), in-degree centrality 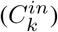, out-degree centrality 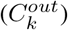, and shortest path length (*SPL*). The energy was defined as the sum of the four KS statistics:

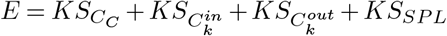

Rather than taking the maximum as in the original formulation (50), we used the sum to ensure that candidate models are evaluated against multiple complementary topological properties simultaneously. Since each of the four KS statistics is bounded in [0, 1], their sum *E* has a maximum of 4 (all four distributions completely non-overlapping) and a minimum of 0 (perfect match on all four).

#### Early stage: Edge addition models

For all edge addition models, the number of edges were added to match the density in the empirical graph. Three edge addition mechanisms were evaluated as candidate wiring rules for the growth phase:

- **Random addition**. Edges were added uniformly at random between pairs of unconnected nodes, serving as a null model.
- **Preferential attachment** (PA). New edges were added via a source-preferential attachment mechanism. The source node *v* was selected with probability proportional to 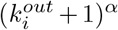, where 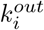 is its out-degree and *α >* 1 introduces super-linear attachment bias. The target node *j* was selected uniformly at random from nodes not yet connected to *i*. With probability *P*_*r*_, an existing edge was removed instead of adding a new one; removed edges were selected via a node chosen with probability proportional to (*k*_*i*_ + 1)^−*γ*^, biasing removal toward low-degree nodes.
- **Preferential attachment with triad formation (PA+TF)**. Following each PA step, with probability *P*_*tf*_ an additional edge was added from *i* to a neighbour of *j* (either a successor or predecessor), promoting local clustering. This mechanism is adapted from the Holme–Kim model (104).

#### Late stage: Node removal models

To model the late-stage decline in network size, we introduced an iterative two-phase model alternating between node removal and edge rewiring. The number of nodes to remove was determined directly from the data as Δ*N* = *N*_0_ −*N*_exp_, where *N*_0_ and *N*_exp_ are the node counts of the baseline and target graphs respectively. Networks exhibiting a node count increase were excluded from the node removal analysis, which occurred in one network of patient C2.

At each iteration, a node was removed with probability *P*_*d*_, or an edge was added via PA+TF with probability 1 −

*P*_*d*_, until all Δ*N* nodes had been removed. Once node removal was complete, edges were added via PA+TF to match the target edge density on the pruned graph, computed as *E*_target_ = *ρ*_exp_ *N*_pruned_(*N*_pruned_ −1), where *ρ*_exp_ is the density of the experimental graph.

Six node removal mechanisms were evaluated, differing in how the node to be removed was selected at each removal step:

- **Random**. Each node was equally likely to be removed, serving as a null model.
- **In-degree**. Nodes were selected for removal with probability proportional to 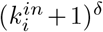, biasing removal toward nodes that receive many connections.
- **Out-degree**. Nodes were selected with probability proportional to 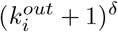, biasing removal toward nodes that send many connections.
- **Degree**. Nodes were selected with probability proportional to (*k*_*i*_ + 1)^*δ*^, biasing removal toward the most connected hubs overall.
- **Clustering high**. Nodes were selected with probability proportional to (*C*_*C,i*_ + *ϵ*)^*δ*^, biasing removal toward nodes deeply embedded in local cliques, consistent with excitotoxicity in overactive local circuits.

**Clustering low**. Nodes were selected with probability proportional to (1 −*C*_*C,i*_ +*ϵ*)^*δ*^, biasing removal toward weakly integrated nodes, consistent with a “use it or lose it” pruning principle.

In all degree- and clustering-based mechanisms, *δ* ≥ 0 controls the nonlinearity of the selection bias: *δ* = 0 recovers uniform random removal, *δ* = 1 gives linear bias, and *δ >* 1 gives super-linear bias. The random removal mechanism has no selection bias and *δ* was therefore not fitted for this condition.

#### Parameter fitting and cross-validation

Model parameters were optimised using Bayesian hyperparameter search (Optuna, 50 trials per fold) to minimise the mean energy across training graphs (105). To prevent overfitting, we employed a stratified network-level cross-validation scheme. Each of the three chips per patient contributed nine networks. In each fold, one network per chip was held out for validation. Parameters were fitted on the remaining networks and evaluated on the three held-out networks.

For the early-stage growth models (PA and PA+TF), the free parameters were *α* ∈ [1.5, 2.0], *γ* ∈ [0, 1], *P*_*r*_ ∈ [0.1, 0.3], and (for PA+TF only) *P*_*tf*_ ∈ [0.5, 1.0]. For the late-stage node removal models, growth-phase parameters were fixed to the mean best-fitting PA+TF parameters from the early-stage analysis, and only the selection strength *δ* ∈ [0, 2] and the node removal probability *P*_*d*_ ∈ [0.1, 1.0] were optimised per removal mechanism. Model comparison was performed on held-out validation energies.

## Random forest models

We used random forest (RF) regression to quantify how much of each population burst feature could be predicted from the static structure of the effective network. For every target, we trained a separate RF on the full set of structural features, *p* in total, estimated its out-of-sample performance by cross-validated *R*^2^, and assessed significance against a label-shuffled null. All forests used 200 trees with 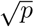 features considered per split and a minimum of 2 samples per leaf; squared error was used as the splitting criterion, and all remaining hyperparameters were left at their SCIKIT-LEARN (v1.7.1) defaults.

We trained one model per burst feature. The targets were the network burst rate, mean network burst duration, the coefficient of variation of the inter-burst interval, within-burst spike rate, burst fragmentation, mean firing rate per neuron, mean pairwise spike-count correlation (rCC), and the percentage of active electrodes. All targets except the percentage of active electrodes were log(1 + *x*)-transformed before fitting to reduce their heavy right skew.

Each model used the same structural descriptors of the effective network as inputs, spanning connectivity, degree-distribution shape, clustering, centrality, path structure, and community and component organisation (Table 2). Extensive quantities that scale with network size were normalised by the number of nodes.

**Table 1.** Model parameters.

| Symbol | Description | Value or range |
| --- | --- | --- |
| $C$ | Membrane capacitance | 281 pF |
| $g_L$ | Leak conductance | 30 nS |
| $E_L$ | Leak reversal / resting | -70.6 mV |
| $V_T$ | Exponential threshold (spike) | -50.4 mV |
| $\Delta_T$ | Slope factor | 2 mV |
| $a$ | Subthreshold adaptation | 4 nS |
| $b$ | Spike-triggered adaptation increment | 0.05 nA |
| $V_{\text{reset}}$ | Reset potential | -62 mV |
| $\tau_w$ | Adaptation time constant | 150 ms ($\pm 30\%$ per neuron) |
| $t_{\text{ref}}$ | Refractory period | 2 ms |
| $E_{\text{exc}}$ | Excitatory reversal | 0 mV |
| $\tau_{\text{rise}}$ | Synaptic rise | 0.1 ms |
| $\tau_{\text{decay}}$ | Synaptic decay | 2 ms |
| $\tau_{\text{poi}}$ | Poisson current decay | 5 ms |
| $K$ | Poisson trains per neuron | 50 |
| $Q$ | Poisson event amplitude | 0.1 nA |
| $N$ | Neurons per network | 60 |
| $\Delta t$ | Integration step | 0.1 ms |
| $T$ | Simulation duration | 125 s |
|  | Discarded transient | 5 s |
| $w$ | Mean synaptic conductance | [5, 200] nS |
| $\nu$ | Poisson rate | [1, 10] Hz |

**Table 2.** Structural features used as inputs to the random forest models. Features marked † were normalised by the number of nodes.

| Family | Feature |
| --- | --- |
| Connectivity | Density |
|  | Reciprocity |
| Degree structure | Standard deviation of in/out-degree$^\dagger$ |
|  | Skewness of in/out-degree |
|  | Kurtosis of in/out-degree |
|  | In-in, out-out, in-out assortativity |
| Clustering | Mean/Standard deviation local clustering coefficient |
|  | Transitivity |
| Centrality | Mean/Standard deviation in-closeness |
|  | Mean/Standard deviation out-closeness |
|  | Mean/Standard deviation node betweenness |
| Path structure | Mean/Standard deviation/max shortest path length |
|  | Global efficiency |
| Mesoscale organisation | Modularity |
|  | Number of communities |
| | Largest-community size$^\dagger$ |
| | Number of Strongly Connected Components$^\dagger$ |
|  | Variance of Strongly Connected Components size |
| | Maximum Strongly Connected Components size$^\dagger$ |

**Table 3.** Functional feature names and abbreviations used throughout the analysis, grouped by category.

| Category | Subcategory | Feature name | Abbreviation | Units |
| --- | --- | --- | --- | --- |
| *Neuron-level functional features* |  |  |  |  |
| | | Firing rate | $FR$ | Hz |
| | | Firing rate inside network burst | $FR_{in}$ | Hz |
| | | Firing rate outside network burst | $FR_{out}$ | Hz |
| | | Mean first spike latency | $FSL$ | s |
| *Network-level functional features* |  |  |  |  |
| | | Network burst rate | $NBR$ | bursts/min |
| | | Coefficient of variation of inter-network burst interval | $CV(NIBI)$ | — |
| | | Network burst duration | $NBD$ | s |
| | | Network burst spike rate | $NBSR$ | Hz |
| | | Percentage of neurons active in network burst | $Pct_{active}$ | % |
| | | Fragments in network bursts | $N_{frag}$ | — |
| | | Correlation coefficient (instantaneous) | $rCC$ | — |

**Table 4.** Structural feature names and abbreviations used throughout the analysis, grouped by category.

| Category | Subcategory | Feature name | Abbreviation |
| --- | --- | --- | --- |
| *Neuron-level structural features* |  |  |  |
| | | Clustering coefficient (node) | $C_c$ |
| | | In-degree centrality | $C_k^{in}$ |
| | | Out-degree centrality | $C_k^{out}$ |
| | | Node betweenness centrality | $C^{bw}$ |
| | | In-closeness centrality | $C_c^{in}$ |
| | | Out-closeness centrality | $C_c^{out}$ |
| | | Z-score broadcasting hubs | $Z_{BH}$ |
| | | Z-score processing hubs | $Z_{PH}$ |
| | | Z-score communication hubs | $Z_{CH}$ |
| *Network-level structural features* |  |  |  |
| | Size | Number of nodes | $N$ |
| | Size | Number of edges | $E$ |
| | Size | Density | $D$ |
| | Clustering | Mean clustering coefficient | $\mu(C_c)$ |
| | Clustering | Standard deviation of clustering coefficient | $\sigma(C_c)$ |
| | Clustering | Transitivity | $T$ |
| | Clustering | Reciprocity | $R$ |
| | Centrality | Mean node betweenness | $\mu(C^{bw})$ |
| | Centrality | Standard deviation of node betweenness | $\sigma(C^{bw})$ |
| | Centrality | Mean in-closeness centrality | $\mu(C_c^{in})$ |
| | Centrality | Standard deviation of in-closeness centrality | $\sigma(C_c^{in})$ |
| | Centrality | Mean out-closeness centrality | $\mu(C_c^{out})$ |
| | Centrality | Standard deviation of out-closeness centrality | $\sigma(C_c^{out})$ |
| | Centrality | Standard deviation of in-degree centrality | $\sigma(C_k^{in})$ |
| | Centrality | Skewness of in-degree centrality distribution | $Skew[C_k^{in}]$ |
| | Centrality | Kurtosis of in-degree centrality distribution | $Kurt[C_k^{in}]$ |
| | Centrality | Standard deviation of out-degree centrality | $\sigma(C_k^{out})$ |
| | Centrality | Skewness of out-degree centrality distribution | $Skew[C_k^{out}]$ |
| | Centrality | Kurtosis of out-degree centrality distribution | $Kurt[C_k^{out}]$ |
| | Paths | Mean shortest path length | $SPL$ |
| | Paths | Standard deviation of shortest path length | $\sigma(SPL)$ |
| | Paths | Maximum shortest path length (diameter) | $max(SPL)$ |
| | Paths | Global efficiency | $Eff$ |
| | Communities | Number of strongly connected components | $SCC$ |
| | Communities | Variance in strongly connected component size | $Var[N_{SCC}]$ |
| | Communities | Fraction of nodes in largest strongly connected component | $f(N_{SCC}^{max})$ |
| | Communities | Modularity | $Q$ |
| | Communities | Number of communities | $M$ |
| | Communities | Fraction of neurons in largest community | $f_{LC}$ |
| | Assortativity | In-degree – out-degree assortativity | $r^{in-out}$ |
| | Assortativity | In-degree – in-degree assortativity | $r^{in-in}$ |
| | Assortativity | Out-degree – out-degree assortativity | $r^{out-out}$ |

**Table 5.** Primary and secondary antibodies used for immunocytochemistry.

| **Antibody** | **Target/Specificity** | **Host** | **Supplier** |
| --- | --- | --- | --- |
| *Primary antibodies* |  |  |  |
| Phalloidin | F-actin | n.a. | Abcam AB176757 |
| Synapsin I mAb | Synapsin I | Rabbit | Abcam, AB254349 |
| PSD-95 Antibody (6G6-1C9) | PSD95 | Mouse | Invitrogen, MA1-045 |
| GFAP pAb | GFAP | Chicken | Abcam, AB4674 |
| Ki67 (SP6) mAb | Ki67 | Rabbit | Abcam, AB16667 |
| Nestin | Nestin | Mouse | Abcam, AB6320 |
| S100β | S100β | Rabbit | Abcam, AB52642 |
| SOX9 | SOX9 | Rabbit | Abcam, AB185966 |
| *Secondary antibodies* |  |  |  |
| Goat anti-mouse Alexa Fluorophore 488 | Anti-rabbit | Goat | Invitrogen A11008 |
| Goat anti-mouse Alexa Fluorophore 555 | Anti-mouse | Goat | Invitrogen A32927 |
| Goat anti-mouse Alexa Fluorophore 647 | Anti-chicken | Goat | Invitrogen A21449 |

Recordings were restricted to DIV 28 to 60. We fit models on four overlapping subsets of the data: a pooled model across all patients; separate early (DIV 28 to 45) and late (DIV 45 to 60) models; and per-patient models, fit only for patients with at least 30 recordings spanning at least 3 chips.

All cross-validation used grouped splitting with the chip as the grouping unit, ensuring that no chip appeared in both training and test sets. We used repeated group shuffle splits with 20 repeats, holding out 3 chips per split for the pooled, early, and late models and 2 chips per split for the per-patient models, owing to their smaller size. Performance was summarised as the mean cross-validated *R*^2^ across splits. To obtain a null distribution, we repeated the full cross-validation 20 times with the target labels randomly permuted and computed an empirical *p*-value as the fraction of label-shuffled runs whose mean *R*^2^ matched or exceeded the observed value. Models with *p*-value *>* 0.05, or *R*^2^ *<* 0.1 were considered not significant. After evaluation, each model was refit on all available data and retained for SHAP feature-importance analysis.

### Gate sweep and mixture model

To explain the observed shift in PID structure over disease progression, we developed a minimal three-node model in which the PID decomposition of any triplet (*X*_1_, *X*_2_ → *Y*) is fully determined by two parameters: the integration mode of the target neuron *Y* (gate type) and the common drive strength *ρ* of its inputs. The common drive model consists of a hidden upstream node *U* (common drive) that co-drives two source neurons *X*_1_ and *X*_2_, which in turn drive a target neuron *Y* . All variables are binary, representing the presence or absence of a spike in a given time bin. The generative model is:

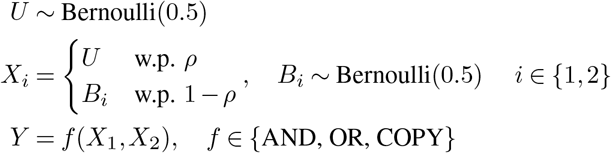

where *ρ* ∈ [0, 1] is the common drive strength, related to the resulting pairwise correlation between *X*_1_ and *X*_2_. At *ρ* = 0, *X*_1_ and *X*_2_ are fully independent. At *ρ* = 1, *X*_1_ = *X*_2_ = *U* always. The gate function *f* determines the integration mode of *Y* : an OR gate (low threshold, fires to any input) or AND gate (high threshold, fires only to coincident inputs) reflects the excitability of the target neuron, and a COPY gate (Y relays one dominant input) reflects sparse, specific connectivity. The full marginal joint distribution *P* (*X*_1_, *X*_2_, *Y*) is obtained analytically by marginalising over *U* :

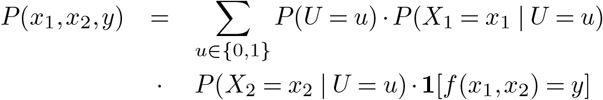

where *X*_1_ and *X*_2_ are conditionally independent given *U* . PID was computed across *ρ* ∈ [0, 1] (60 equally spaced values) for each gate type (AND, OR, COPY), yielding the PID trajectory as a function of common drive strength. The PID of each triplet was computed using the BROJA estimator (94) as described in “BROJA estimator implementation” in the Supplementary Material. Because no single gate type can simultaneously reproduce the observed symmetric increase in both redundancy and synergy, we modelled the network as a mixture of two populations: COPY-like triplets (weight *w*_copy_, reflecting sparse specific connectivity) and integrative triplets (weight 1− *w*_copy_, reflecting a 50/50 mixture of OR and AND gates at common drive strength *ρ*). The predicted PID fractions are:

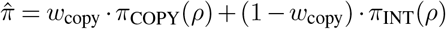

where *π* = (Red, Unq, Syn) are normalised fractions of total MI, *π*_COPY_(*ρ*) is the COPY-like gate fingerprint at common drive strength *ρ* and *π*_INT_(*ρ*) is the integrative gate fingerprint at common drive strength *ρ*.

#### Solution space

For each *ρ*, the optimal *w*_copy_ was obtained analytically via least squares:

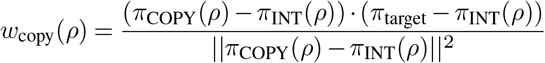

clipped to [0, 1]. The pair (*w*_copy_, *ρ*) minimising the squared residual was selected as the optimal solution. Each condition and timepoint was fitted independently, allowing *ρ* to vary across the disease progression.

### State space quantification

Starting from the binary spike matrix, spikes were binned into 20 ms windows, yielding a coarser activity matrix in which each column represents the spatial pattern of active neurons during one time bin, referred to here as a population state (54, 55, 106). A bin width of 20 ms was chosen to match the typical timescale of synaptic integration in cortical networks, following established practice in population state analyses (54, 55). We enumerated all unique population states across the recording and computed their empirical probability distribution, from which Shannon entropy was calculated as:

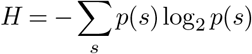

To account for differences in the number of observed states across recordings, entropy was normalised as *H/H*_max_, where *H*_max_ = log2(*S*_unique_), yielding a value of 1 for a maximally uniform state distribution and values approaching 0for stereotyped or repetitive dynamics. The number of unique states normalised by neuron count (*S*_unique_*/N*) was additionally computed as a size-independent measure of state space richness.

### Quantification and statistical analysis

All statistical analyses were performed in Python. The sample size refers to the number of networks or individual nodes as specified in each figure legend. In total, we recorded 60 Neuro-Astro, 52 Neuron-only, 26 Neuro-GBM C1, 27 Neuro-GBM C2, and 27 Neuro-GBM M1 networks. For comparisons across three or more groups, a Kruskal–Wallis omnibus test was performed, followed by Dunn’s post-hoc test with Benjamini–Hochberg correction for multiple comparisons. Statistical significance is denoted as: ns (p > 0.05), * (0.01 < p ≤ 0.05), ** (0.001 < p ≤ 0.01), *** (0.0001 < p ≤ 0.001), and **** (p ≤ 0.0001). For longitudinal line plots tracking network trajectories over days *in vitro*, data were grouped into bins of 5 DIVs to account for minor variation in recording timepoints and fluctuations in culture medium composition. Where multiple recordings fell within the same bin, their mean was used as the statistical unit. Box plots show median, interquartile range, and 1.5× IQR whiskers. Bar plots show the mean ± standard deviation or mean ± SEM as indicated in the figure legend. Detailed statistical results for all comparisons are provided with the published data.

### Ethics statement

Primary glioblastoma cells were derived from tumour tissue resected from patients undergoing surgery at the University Hospital Basel, Switzerland. All donors provided written general consent for the further use of their biological material for research purposes. Use of the material was approved by the Ethics Committee of Northwestern and Central Switzerland (EKNZ; BASEC No. 2024-00218). All experiments were performed in accordance with the Declaration of Helsinki and the Swiss Human Research Act (HRA).

## SUPPLEMENTARY INFORMATION

### BROJA estimator implementation

The PID of each triplet was computed using the BROJA estimator (94). The BROJA redundancy is defined as:

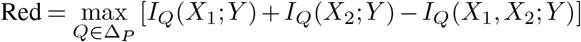

where Δ_*P*_ = {*Q*(*X*_1_, *X*_2_, *Y*) : *Q*(*X*_1_, *Y*) = *P* (*X*_1_, *Y*) and *Q*(*X*_2_, *Y*) = *P* (*X*_2_, *Y*)} is the set of distributions preserving the bivariate marginals of the observed distribution *P* and *I*_*Q*_ is the mutual information evaluated on the distribution *Q*. Since the constraints fix *I*_*Q*_(*X*_1_; *Y*) = *I*_*P*_ (*X*_1_; *Y*) and *I*_*Q*_(*X*_2_; *Y*) = *I*_*P*_ (*X*_2_; *Y*) for all *Q* ∈ Δ_*P*_ , maximising the redundancy reduces to minimising *I*_*Q*_(*X*_1_, *X*_2_; *Y*) over Δ_*P*_.

The optimisation was parametrised via the conditional distribu tion 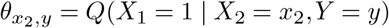 , which yields four free parameters for binary variables. The equality constraints 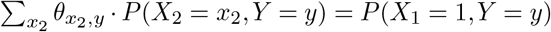 enforce preservation of the *P* (*X*_1_, *Y*) marginal. The constrained optimisation was solved using Sequential Least Squares Programming (SLSQP) with *n* = 50 random restarts for numerical stability, implemented in Python using `scipy.optimize.minimize`. Bounds were tightened analytically when constraints forced parameters to boundary values, improving convergence near degenerate distributions. The remaining PID components follow from the lattice identities:

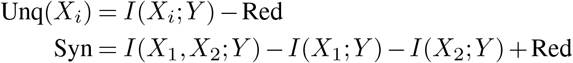

The implementation was verified against known analytical results: XOR yields pure synergy (Syn = 1, Red = Unq = 0) and COPY yields pure unique information (Unq(*X*_1_) = 1, Red = Syn = 0).

### Ground-truth simulation model

Each node was an adaptive exponential integrate-and-fire (AdEx) neuron (44) with membrane potential *v* and adaptation current *w*:

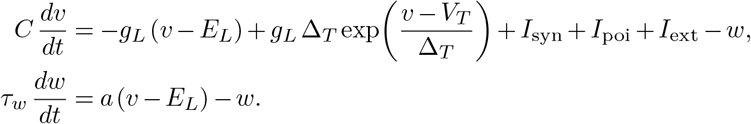

A spike was registered when *v* crossed the exponential threshold *V*_*T*_ , upon which *v* ← *V*_reset_ and *w* ← *w* + *b*, followed by a 2 ms refractory period. Subthreshold parameters (*C, g*_*L*_, *E*_*L*_, *V*_*T*_ , Δ_*T*_ , *a, b, V*_reset_) followed the AdEx regular-spiking set (44). *τ*_*w*_ carried per-neuron heterogeneity (± 30% uniform jitter).

Two independent sources drove each neuron. (i) A constant bias current 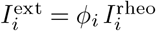, where 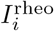 is the AdEx rheobase computed analytically from neuron *i*’s parameters and *ϕ*_*i*_ ∼ *U* [0, 0.3]. (ii) An excitatory shot-noise background modelled as a decaying current 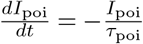 , receiving *K* = 50 independent homogeneous Poisson trains per neuron, each of rate ν, with every event adding *Q* = 0.1 nA to *I*_poi_.

Synapses are modelled as conductance-based and excitatory,

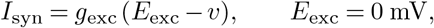

with a bi-exponential time course generated by a rise and a decay variable,

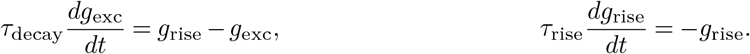

Networks were integrated with forward Euler at Δ*t* = 0.1 ms for 125 s in Brian2 (95); the first 5 s were discarded as transient, leaving 120 s for analysis.

**Fig. S1.**
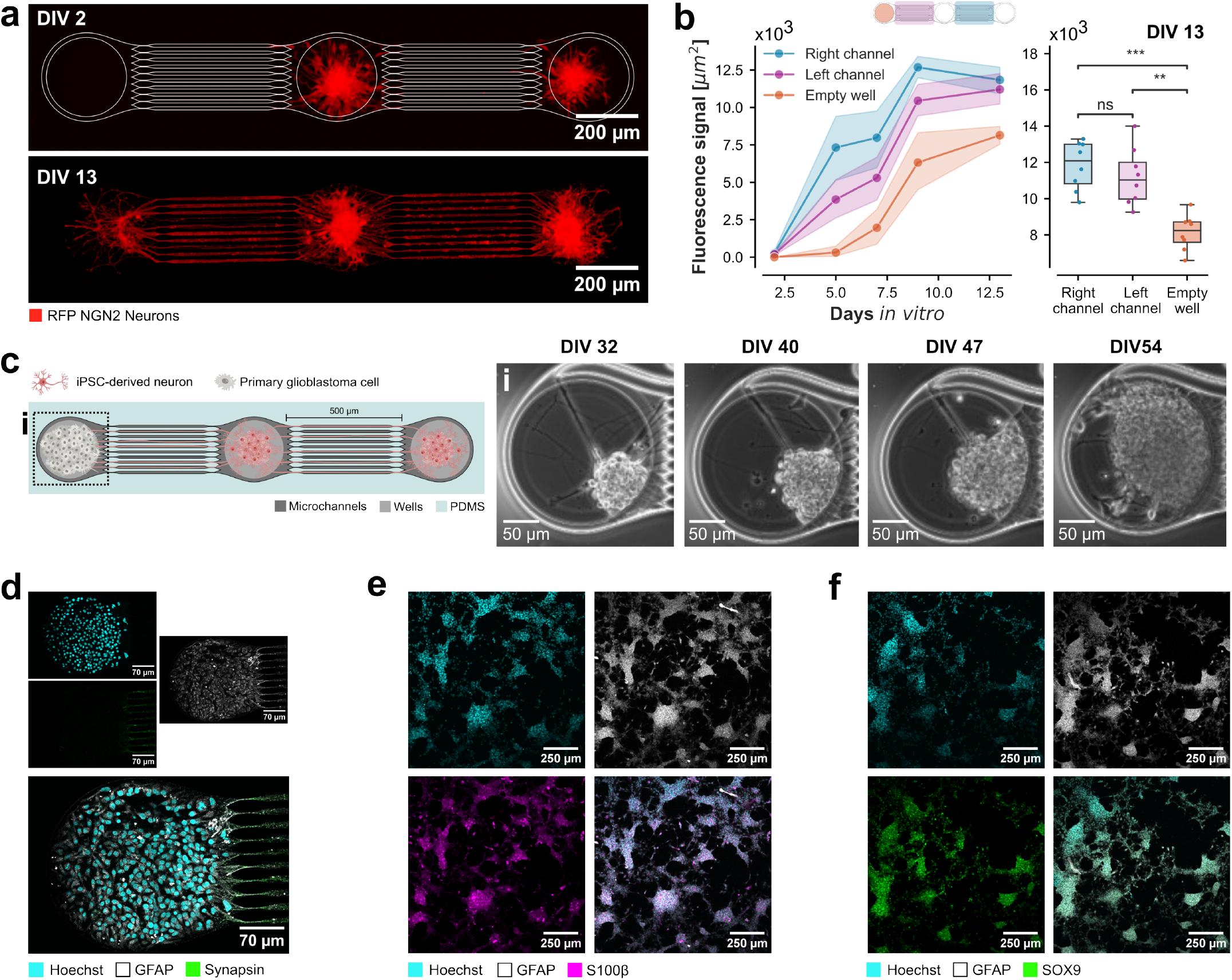
Additional characterisation of the *in vitro* co-culture platform. **a** Representative fluorescence images showing neurite growth at DIV 2 (top) and DIV 13 (bottom). **b** Fluorescence signal representing neurite presence over days *in vitro* (left) and distribution at DIV 13 (right). Lines denote the mean and shaded bands the bootstrapped 95 % confidence interval across networks. Boxplot shows median and inter-quartile range, with individual ROIs shown as dots. No significant difference in signal was detected between the two channel sides after two weeks *in vitro* (Kruskal–Wallis with Dunn’s post-hoc and Benjamini–Hochberg correction, ns, not significant, p > 0.05; *p < 0.05; **p < 0.01; ***p < 0.001; ****p < 0.0001). **c** Glioblastoma spheroids remained spatially confined to the third seeding well throughout the co-culture period despite ongoing proliferation. Representative brightfield images are shown at DIV 32, DIV 40, DIV 47, and DIV 54. **d** Representative image of iPSC-derived astrocytes in the third seeding well of positive control conditions at DIV 35. GFAP staining highlights astrocytes; Synapsin staining labels neuronal neurites. **e–f** Astrocytes maintained expression of the glial markers S100*β*, SOX9, and GFAP in co-culture medium.

**Fig. S2.**
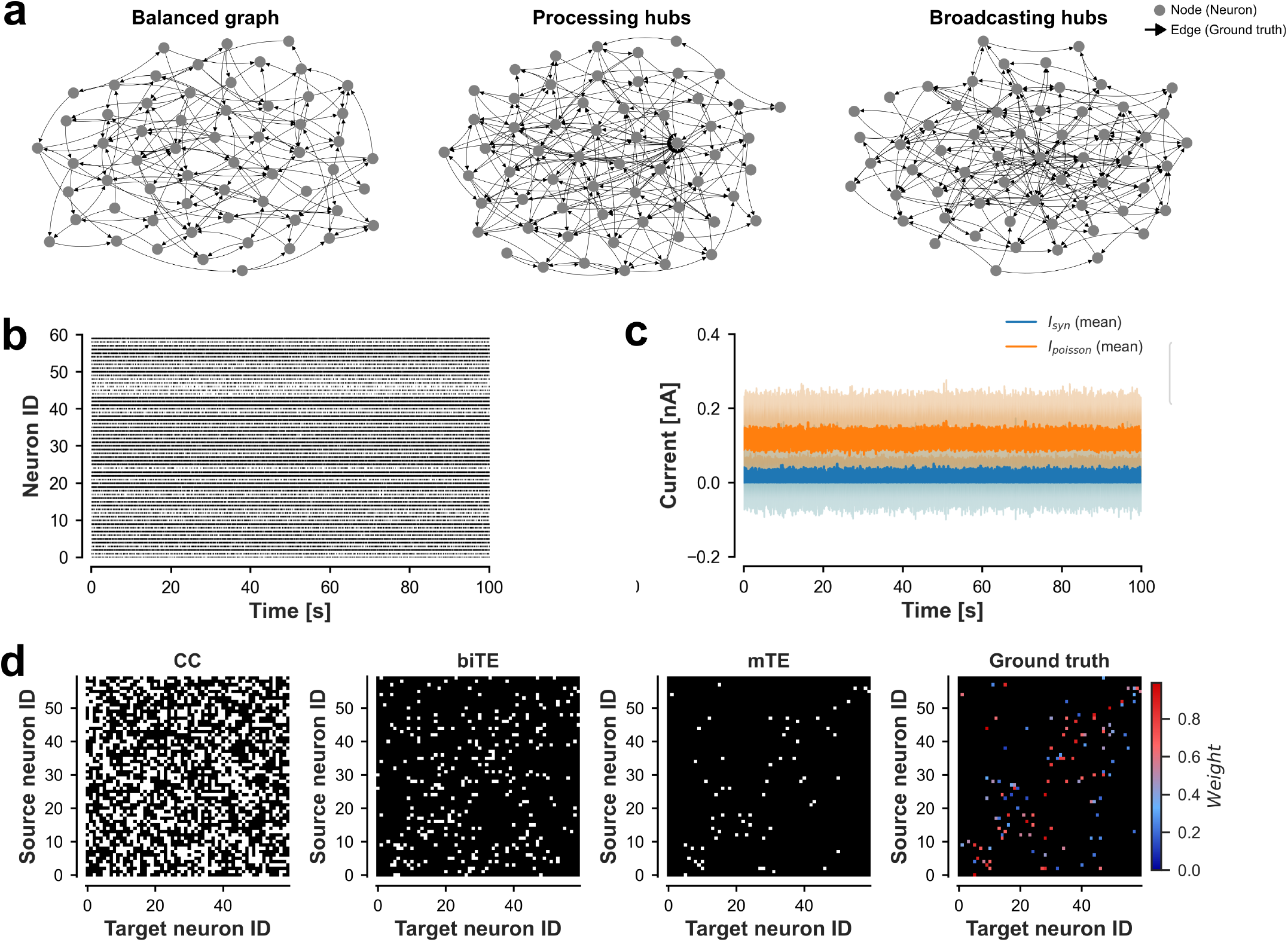
Assessment of multivariate transfer entropy (mTE)-based network inference using simulated networks. **a** Representative examples of the three generated ground-truth network topologies. Starting from an Erdő s–Rényi baseline, three randomly selected nodes were assigned additional outgoing (broadcasting hub class) or incoming (processing hub class) edges. **b** Simulated spike trains from one such network, together with the corresponding **c** synaptic and Poisson input-current traces. Mean synaptic current was consistently lower than mean Poisson current. **d** Adjacency matrices inferred from the spike trains in **b** using cross-correlation (CC), bivariate transfer entropy (biTE), and mTE.

**Fig. S3.**
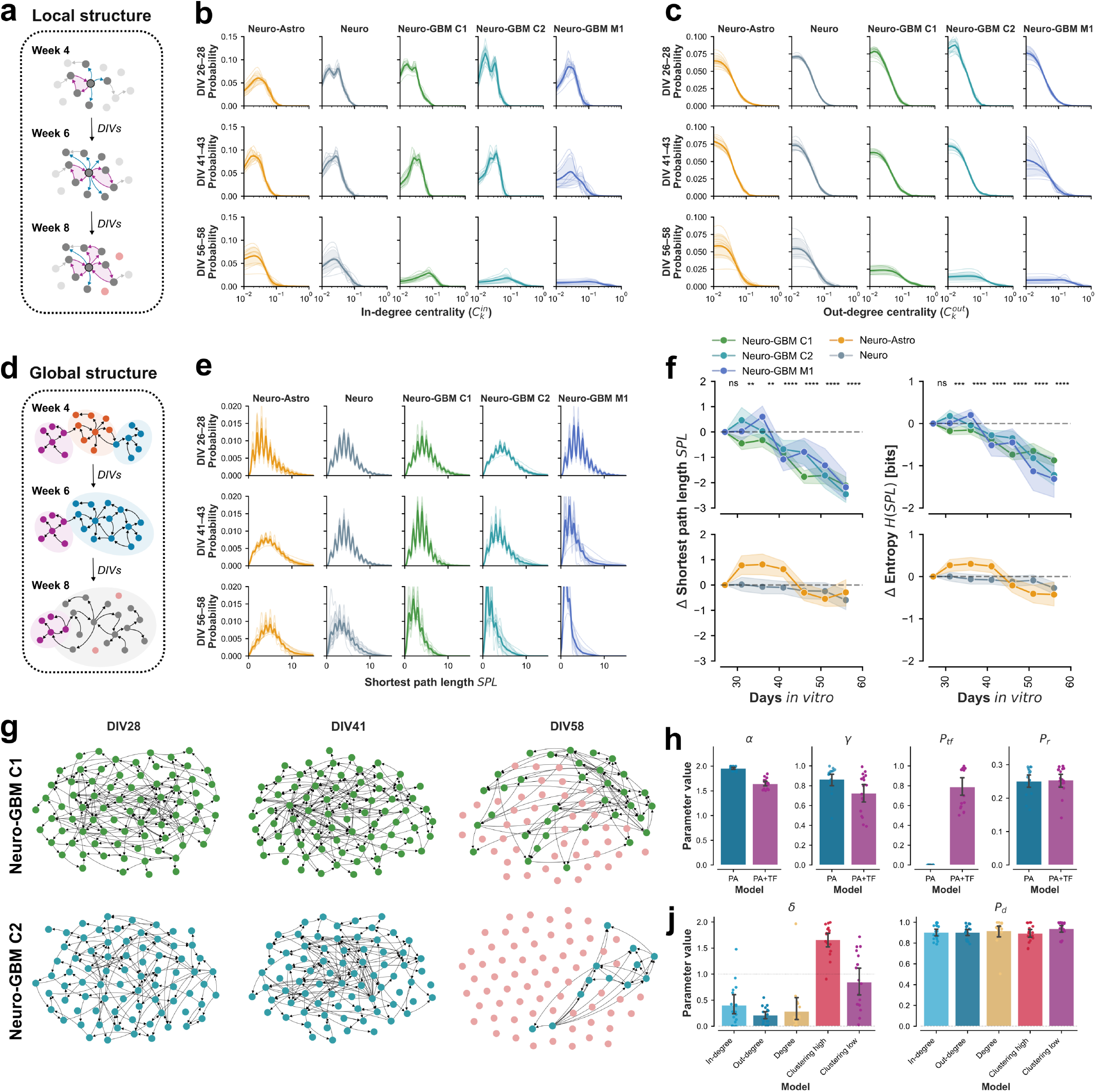
Glioblastoma invasion progressively restructures both local and global network topology. **a** Schematic illustrating changes in local network structure across Weeks 4, 6, and 8 *in vitro*, highlighting redistribution of connectivity. **b–c** Probability distributions of in-degree centrality 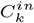 and out-degree centrality 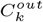 across all five conditions at DIV 26–28, DIV 41–43, and DIV 56–58. Transparent lines show individual networks’ KDE. Saturated lines show the mean and standard deviation across networks. **d** Schematics illustrating changes in global network structure over the same period, showing progressive fragmentation and community reorganisation. **e** Probability distributions of shortest path length *SPL* across conditions and timepoints, formatted as in **b–c. f** Change in mean shortest path length Δ*SPL* and change in *SPL* entropy Δ*H*(*SPL*) relative to baseline, plotted over days *in vitro*. Δ*H*(*SPL*) was computed in bits as *H*(*SPL*) = −Σ _*l*_ *p*(*l*) log_2_ *p*(*l*), where *p*(*l*) is the fraction of node pairs connected by a path of length *l*. Lines show the mean and shaded bands the bootstrapped 95% confidence interval across networks. Asterisks indicate significant differences between conditions at each timepoint (Kruskal–Wallis with Dunn’s post-hoc and Benjamini–Hochberg correction, ns, not significant, p > 0.05; *p< 0.05; **p < 0.01; ***p < 0.001; ****p < 0.0001). **g** Representative effective connectivity networks for Neuro–GBM C1 (top) and Neuro–GBM C2 (bottom) at DIV 28, DIV 41, and DIV 58 used to train the generative model, illustrating progressive loss of connected neurons (green/teal) and accumulation of isolated nodes (pink) over time. **h** Estimated parameters *α, γ, P*_*tf*_ , and *P*_*r*_ of the preferential attachment (PA) and preferential attachment with topological features (PA+TF) network growth models. **j** Estimated parameters *δ* and *P*_*d*_ of PA+TF model variants with node deletion based on structural features (in-degree, out-degree, degree, clustering high, clustering low). Bars show mean, error bars 95 % CI; individual networks shown as scatter points.

**Fig. S4.**
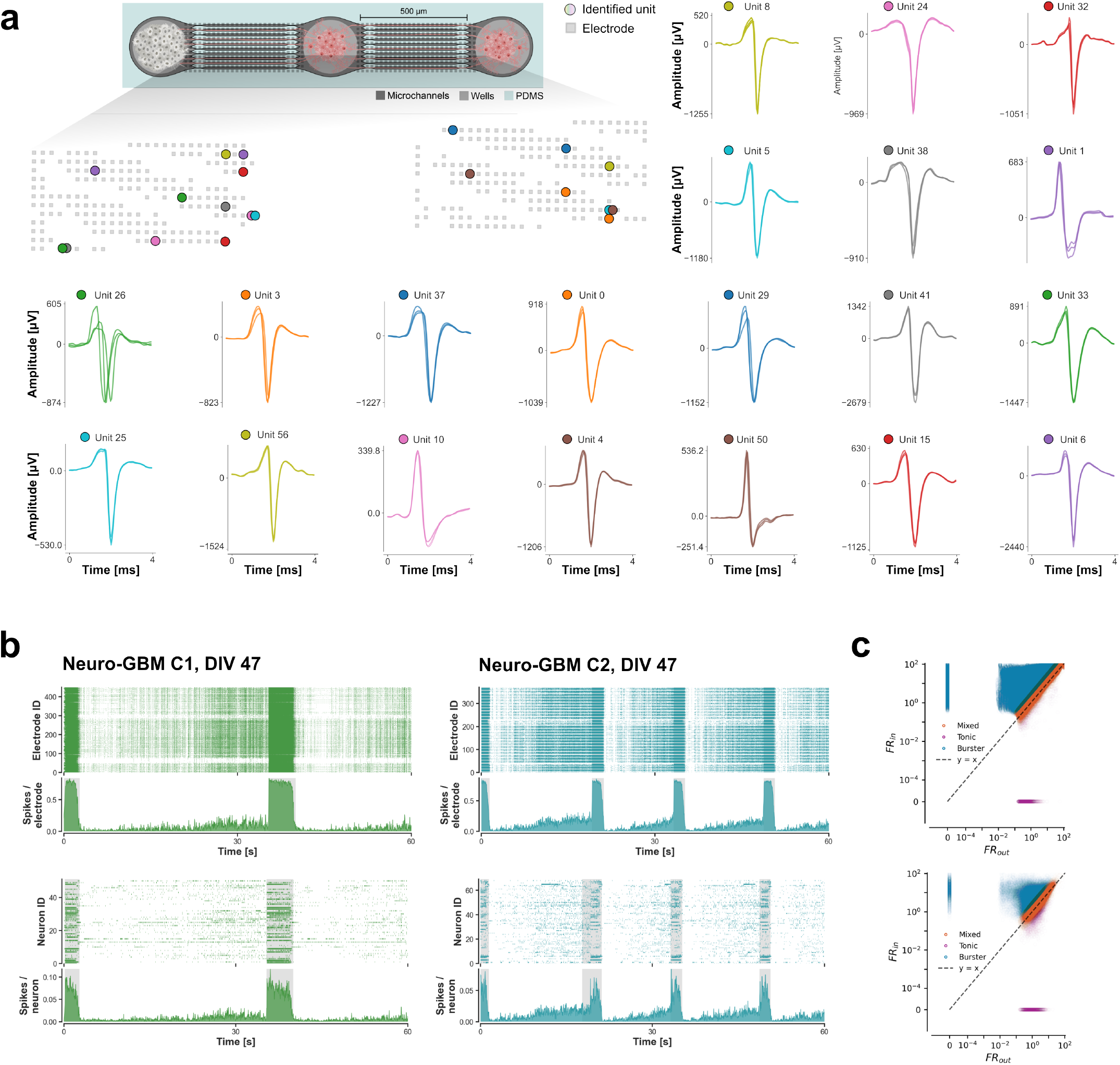
Spike sorting identifies individual neurons and small neuronal clusters from extracellular electrophysiological recordings. **a** Representative spike sorting output obtained using SpikeInterface (85) with SpyKING CIRCUS 2 (86). Waveforms of 20 randomly selected units are plotted alongside the location of the highest-amplitude electrode for each unit on the electrode grid. Waveforms recorded on three electrodes assigned to each unit are shown. **b** Representative raster plots of spike times and population firing rates for unsorted data (per electrode, top) and spike-sorted data (per unit, bottom). Network bursts are indicated by shaded grey regions. Comparable network dynamics are observed across conditions. **c** Classification of burster, mixed, and tonic neurons identified in unsorted (top) and spike-sorted (bottom) recordings. Each neuron is characterised by its mean firing rate inside bursts *FR*_*in*_ relative to its rate outside bursts *FR*_*out*_. Scatter plot shows *FR*_*in*_ versus *FR*_*out*_ across all neurons (DIV 30–40), coloured by firing class; dashed line indicates the identity line.

**Fig. S5.**
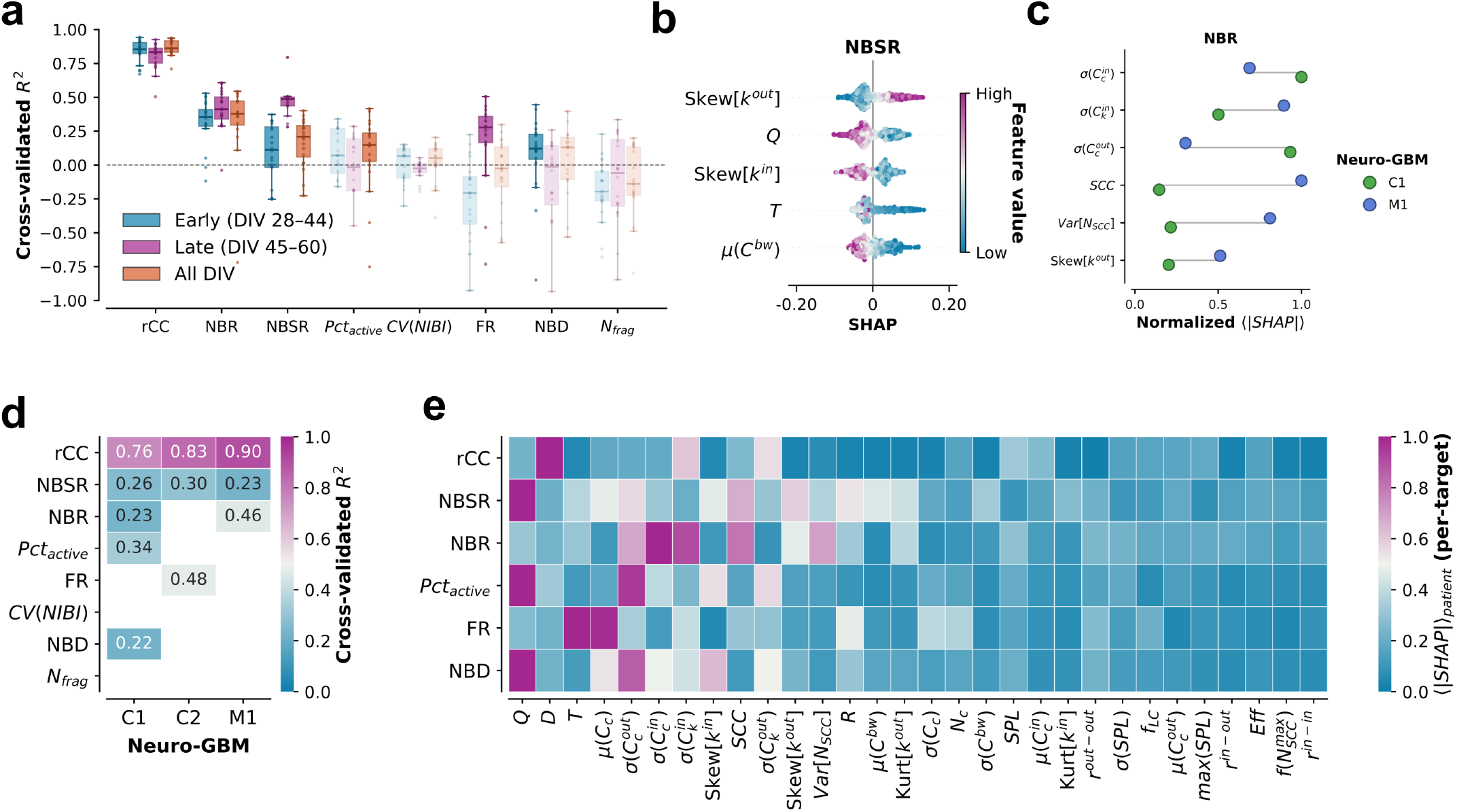
Random forest models of structure-to-dynamics prediction: training regimes, per-patient generalisation, and full feature importances. **a)** Cross-validated *R*^*2*^ for predicting each burst feature from network structure, compared across three training regimes: early (DIV 28 to 44), late (DIV 45 to 60), and all DIV pooled. Boxes show the distribution across cross-validation splits (individual splits overlaid as points); dashed line, *R*^*2*^= 0. *rCC* is predicted well in every regime, *NBR* and *NBSR* moderately, and the remaining features poorly. **b)** SHAP summary (beeswarm) for the within-burst spike rate (*NBSR*) model (top 5 features), pooled across cultures. Each point is one network; horizontal position gives the feature’s signed contribution and colour encodes the feature value (low to high). **c)** Normalised mean absolute SHAP value of the top features for the network burst rate (*NBR*) model, shown per patient; only patients with a significant *NBR* model are displayed (C1, M1). **d)** Cross-validated *R*^*2*^ for predicting every burst feature within each patient line (C1, C2, M1). Only cells with a significant model are shown (*R*^*2*^ *>* 0.05 and permutation *p <* 0.05); blank cells did not reach significance. **e)** Per-target normalised mean absolute SHAP value across the full structural feature set (columns) for each well-predicted target (rows). Values are averaged over patients and scaled to each target’s maximum (per-target normalisation); non-significant (patient, target) models were excluded before averaging.

**Fig. S6.**
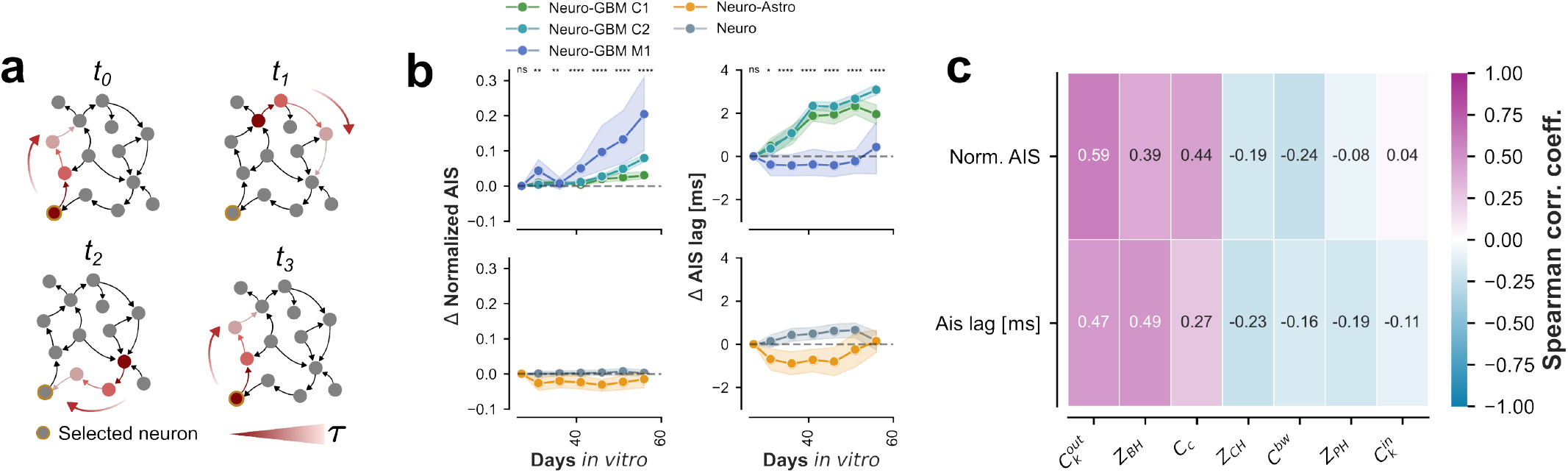
Glioblastoma co-culture increases active information storage within neuronal clusters. **a** Schematic illustrating the concept of active information storage (AIS), which quantifies the degree to which a neuron’s current activity can be predicted from its own past activity. This can be interpreted as the extent to which previously stored information is reactivated within local neuronal clusters (60). **b** Change in normalised AIS and AIS lag relative to baseline over days *in vitro*. Normalised AIS increases disproportionately in patient-derived conditions (Neuro–GBM C1, C2, and M1), while AIS lag increases predominantly in Neuro–GBM C1 and C2. Lines show the mean and shaded bands the bootstrapped 95 % confidence interval across networks. Asterisks indicate significant differences between conditions at each timepoint (Kruskal–Wallis with Dunn’s post-hoc and Benjamini–Hochberg correction, ns, not significant, p > 0.05; *p< 0.05; **p < 0.01; ***p < 0.001; ****p < 0.0001). **c** The mean per-network increase in normalised AIS and AIS lag correlated with out-degree centrality 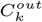, broadcasting hub z-score *Z*_*BH*_ , and clustering coefficient *C*_*C*_ , suggesting that information reverberates for longer within highly clustered subnetworks characterised by high out-degree. Correlations were considered weak (0.1 ≤ |*ρ*| *<* 0.3), moderate (0.3 ≤ |*ρ*| *<* 0.5), or strong (|*ρ*| ≥ 0.5), following (53).

**Fig. S7.**
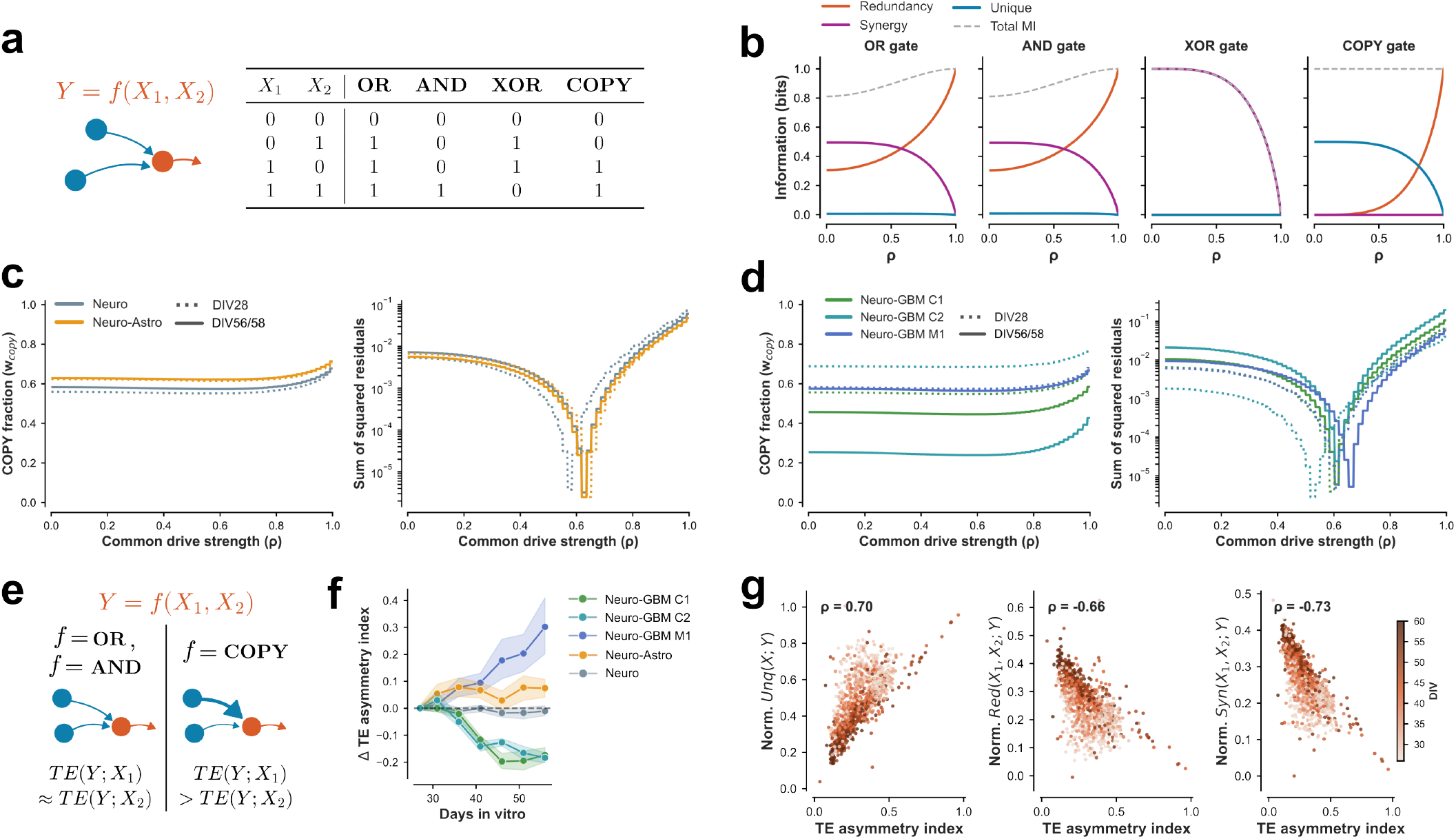
Common drive model links gate-type computation to transfer entropy asymmetry and partial information decomposition components. **a** Truth table for four gate functions (OR, AND, XOR, COPY) commonly used in literature, in which two source neurons *X*_1_ and *X*_2_ determine the firing of target neuron *Y* = *f* (*X*_1_ , *X*_2_). **b** Analytical PID solutions obtained using the BROJA estimator for each gate type, showing redundancy, unique information, synergy, and total mutual information (dashed) as a function of common drive strength *ρ*. OR and AND gates transition from synergy-to redundancy-dominated regimes with increasing *ρ*; the XOR gate remains synergy-dominated throughout; the COPY gate is characterised by high unique information across all *ρ*. **c–d** COPY fraction *w*_*copy*_ (left) and sum of squared residuals (right) as a function of *ρ* for control conditions (Neuro, Neuro–Astro; **c**) and patient-derived GBM conditions (Neuro–GBM C1, C2, M1; **d**) at DIV 28 (dotted) and DIV 56/58 (solid). The minimum of the residual curve identifies the optimal *ρ* for each condition and timepoint. **e** Schematic illustrating the relationship between gate type and transfer entropy (TE) asymmetry. OR- and AND-types should have approximately symmetric TE input, whereas COPY-type gates should have one dominant TE stream, reflecting dominant drive from a single source. **f** Change in mean TE asymmetry index relative to baseline over days *in vitro* across all five conditions. Lines show the mean and shaded bands the bootstrapped 95% confidence interval across networks. **g** Scatter plots showing Spearman correlations between the TE asymmetry index and normalised unique, redundant and synergistic information contribution. Each point represents one network; colour encodes DIV.

